# Fragment Based Active Site Exploration of Urethane Hydrolases Reveals a Diversity of Urethane Binding Modes

**DOI:** 10.64898/2026.07.06.734427

**Authors:** Deniz Bicer, Diana Kochubei, Rosie Graham, Samuel Peña–Díaz, Laura Rotilio, Nikolaj L. Villadsen, Andreas Sommerfeldt, Martin B. Johansen, Alexander Sandahl, Søren S. Thirup, J. Preben Morth, Daniel E. Otzen

## Abstract

Recent advances in the discovery, characterisation, and engineering of urethanases provide new opportunities for the sustainable biocatalytic degradation of polyurethane waste. A mechanistic understanding of enzyme–plastic interactions is essential for structure-based engineering to enhance urethanase activity. However, the extremely complex and hydrophobic nature of polyurethane makes it challenging to elucidate the structural basis of enzyme–plastic interactions. Here, we used a fragment-based approach to characterise the active sites of two novel urethanases with different catalytic scaffolds, employing both a crystallographic fragment-screening (FASE) campaign and soluble fragments of plastic-like analogues that mimic the substrate, transition state, or product. FASE identified new substrate-binding subpockets while interactions of plastic mimetics in the active site provided a mechanistic understanding of the recognition and binding of polyurethane fragments by these subpockets. These results highlight a diversity of binding modes among urethanases toward different polyurethane fragments.

**Synopsis:** Fragment-based active site exploration of urethanases to elucidate polyurethane binding and cleavage

## 1. Introduction

Polyurethane (PU) plastics are an essential part of modern society, forming numerous everyday materials due to their ease of manufacture, robustness and versatility (Rossignolo *et al*., 2024). Despite these assets, the alarming accumulation of PU waste in the ecosystem, exceeding millions of tons, makes a pressing case for recycling strategies in the transition to a sustainable and circular economy (Banik *et al*., 2023). Current methods for recycling PU are mostly based on expensive, open-loop and unsustainable mechano-chemical processing steps that generate by-products, many of which are harmful to the environment, such as carbon dioxide (Polecki *et al*., 2026). Biocatalytic, *i.e.* enzymatic, degradation provides an attractive alternative due to mild conditions and highly specific reactions. There is currently a rapid expansion in the discovery, characterization and engineering of polyurethane degrading enzymes or PURases of typically microbial origin (Branson *et al*., 2023; Chen *et al*.,2025; Rotilio *et al*., 2026; Bendtsen *et al*., 2026). Yet most of these novel PURases only present low-level activity on PU plastics, highlighting the urgent need for engineering campaigns to improve catalytic capacity and thermal stability. In this approach, atomic-level mechanistic understanding of enzyme-ligand interactions in the active site is a critical bottleneck. Commercial PU consists of repeating units of an aromatic isocyanate, commonly made from methylene dianiline (MDA) or toluenediamine (TDA), connected through a polyol chain (*e.g.* ethylene glycol) via a urethane bond (**Fig. 1**). Collectively, the aromatic group and the urethane bond constitute a rigid and hydrophobic segment. In contrast, the polyol chain forms a flexible and hydrophilic segment (Akindoyo *et al*., 2016). The polymeric and highly insoluble nature of PU precludes atomic-level structural analysis of enzymes bound to *bona fide* substrates. A trick is to reduce the challenge by turning to either (a) fragments that mimics short segments of the aromatic moiety of PU or (b) short and soluble PU analogues. This provides an alternative approach to explore both the catalytically active site as well as surrounding regions that take part in the binding and release of substrates and products, respectively.

**Figure 1.**
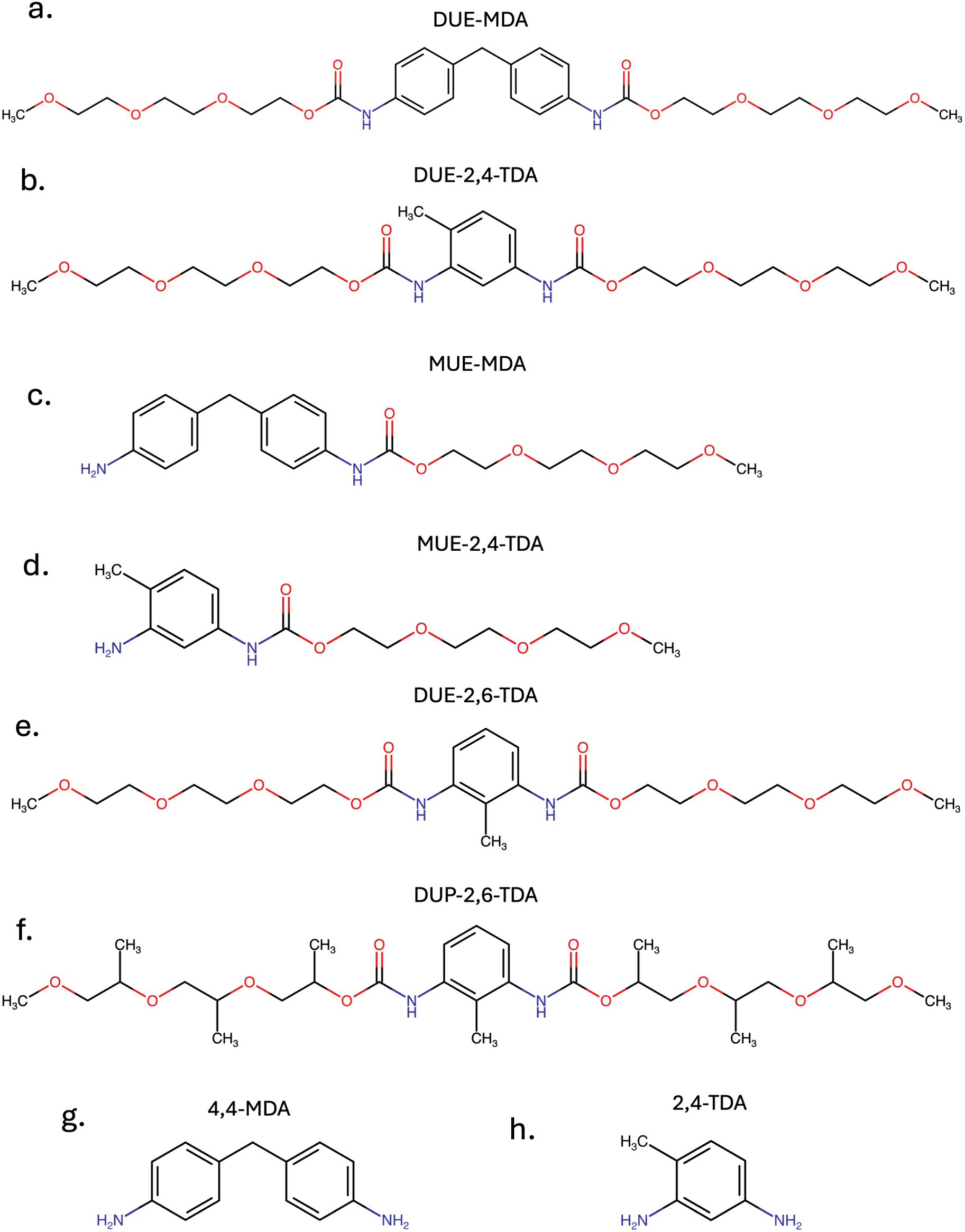
Structures of PU fragments. Substrates: a. DUE-MDA and b.DUE-2,4-TDA. Single urethane bond degraded intermediate products: c. MUE-MDA, d. MUE-2,4-TDA, e)DUE-2,6-TDA, f) DUP-2,6-TDA. Double urethane bond-cleaved products: g. 4,4-MDA and h. 2,4-TDA. DUE, diurethane ethylene glycol; MUE, monourethane ethylene glycol; DUP, diurethane polypropylene glycol. Figure made with *ChemAxon*.

Typical fragments for such studies generally follow the so-called “Rule of Three”, *i.e.* molecular weight < 300 Da, < 3 rotatable bonds and < 3 H-bond donors/acceptors (Jhoti *et al*., 2013). Due to their small size and simplicity, fragments effectively explore chemical space in the binding cavity of an enzyme. For example, in fragment-based drug discovery campaigns, fragments usually serve as the starting point for structure-based drug design (Scott *et al*., 2012). Moreover, fragments can be used to map the binding landscape of an enzyme’s extended active site by binding to all available pockets as potential ligand binding sites. The identification of these binding sites can reveal the molecular basis of enzyme-substrate specificity and hence, establish a rational basis for protein engineering which is critical for most industrially relevant enzymes.

Most currently known urethanases fall into two different serine hydrolase families: i) Amidase Signature (AS) Family Amidases (EC 3.5.1.4) and ii) Carboxylic Acid Esterases (EC 3.1.1.1). These two classes represent different catalytic scaffolds for PU degradation, potentially presenting distinct substrate binding and recognition modes. For this study, we selected two of our recently discovered novel PU-degrading enzymes, namely u11, a member of the Amidase Signature family amidase discovered by database mining (Rotillio *et al*., 2026), and AX^PUR5^, a carboxylic esterase identified from plastic waste landfill (Bendtsen *et al*., 2026). The two enzymes differ in their ability to degrade soluble PU model substrates such as DUE-MDA and DUE-TDA; both enzymes degrade the former, whereas the latter is degraded only by u11.

Here, we employ a small-molecule-based approach to characterise the active-site cavity of PU-degrading enzymes, combining crystallographic fragment screening with binding analysis of PU fragment analogues. Our approach, called Fragment Based Active Site Exploration (FASE), explores the extended active site, *i.e* both the catalytic site and neighbouring binding sites. We first performed a crystallographic fragment screening campaign to probe the active site of enzymes to map subpockets. Guided by these newly mapped pockets, we then analysed the binding of short, soluble plastic fragments that are analogues to the PU substrate, the transition state (TS) or the products of the degradation (**Fig. 1**). The combination of various fragments maps the entire binding landscape for active site of novel PU degrading enzymes and uncovers specific binding poses for PU plastics. Additionally, through FASE, we perform a comparative characterisation of the active sites of u11 and AX^PUR5^ to elucidate PU-binding modes and to explain observed differences in activity across different PU substrates. These insights can guide discovery and engineering efforts of homologous enzymes within same families and provide the molecular basis for the chemical promiscuity inherent to active sites in the AS and esterase family.

## 2. Material and Methods

### 2.1. Expression, Purification and Crystallisation

*Protein expression:* Genes encoding u11/u11-S173A and AX^PUR5^ were cloned into pETM-11 and pET-28 plasmids, respectively, ensuring that each enzyme has a 6xHis tag at the C-terminus, preceded by a TEV cleavage site. The plasmids were transformed into *E.coli* BL21(DE3) super-heat competent cells by heat-shock. Briefly, plasmids were mixed with KCM buffer (0.2 M KCl, 0.2 M CaCl_2_, 0.2M MgCl_2_) on ice for 5 min, then 50 µL of competent cells were added. After further incubation on ice for 20 min, cells were transferred to room temperature and incubated for 5 more min. Then 400 µL 37 ℃ LB media was added, and cells were incubated at 37 °C in a shaking incubator at 150 rpm for 1 h. 100 µL cell suspensions were then dispersed on a LB-Agar plate containing 1 mM Kanamycin and incubated at 37°C overnight, after which a single colony was transferred into 10 mL LB containing 1 mM Kanamycin. This starter culture was grown at 37 ° C overnight with 150 rpm shaking, then transferred to 1 L of LB media supplemented with 1 mM Kanamycin and grown at 37 ℃ at 150 rpm in a shaker until OD600 0.6-0.8, cooled at 4° C for 30 min and induced with 0.5 mM IPTG, after which cells were grown at 18 ℃ at 150 rpm overnight. Cell pellets were harvested at 6000 rpm for 20 min and stored at –20° C until purification.

*Protein purification:* The pellet was suspended in 50 mL Lysis buffer (per L cell culture) containing 50 mM Tris-HCl pH 8.0, 500 mM NaCl, 25 mM Imidazole pH 8.0, 1 mM DTT and 1 mM PMSF (PMSF was omitted for active protein purification), incubated on a rotating table with 60 rpm at 4°C for 30 min and lysed by sonication with 3 cycles of 2 min at 30% power with 1 s pulses. The lysate was cleared by centrifugation for 45 min at 15.000 rpm, after which the supernatant was filtered with a 45 µm membrane filter and loaded onto a HiTRAP 5 mL Ni-NTA column previously washed with x5 CV MQ and then equilibrated with 5X binding buffer (50 mM Tris-HCl pH 8.0, 500 mM NaCl, 25 mM Imidazole pH 8.0) before loading. Protein binding to the column was allowed up to two hours to maximise binding. His-tagged proteins were eluted by elution buffer (50 mM Tris-HCl pH 8.0, 500 mM NaCl, 500 mM imidazole pH 8.0) in stepwise concentration gradients of 10%, 25%, 50% and 100% of the elution buffer. Fractions were run on SDS-PAGE to check purity. To remove the His-tag, fractions with enzymes were dialysed overnight against SEC buffer (50 mM Tris-HCl pH 8.0, 150 mM NaCl) containing TEV protease in 10 kDa cutoff dialysis bag membrane at 4° C with 1 mg TEV protease per 10 mg of enzyme, concentrated in a vivaspin concentrator with cutoff 10 kDa and loaded onto superdex 75 10/300 GL column. Peak fractions were analysed by SDS-PAGE and concentrated with a 10 kDa cutoff to 20 and 40 mg/ml for u11 and AX^PUR5^, respectively (concentrations determined by OD280 using the extinction coefficients for u11 (51700) and AX^PUR5^ (84130)).

Protein crystallisation: Initial crystallisation screening plates were set up in SWISS 3-well plates with different protein:reservoir ratios (1:1, 2:1, and 1:2) at a final drop size of 600 nL and kept at 19 °C. Gradient optimisation of the initial concentrations of salt, precipitants, and protein was conducted in 24-well plates alongside seeding with different dilutions of the initial seed stock.

### 2.2. Fragment soaking and crystal mounting

Plastic fragments including DUE-MDA, MUE-MDA, DUE-2,4-TDA and MUE-2,4-TDA were synthesised as a powder or oil as described elsewhere (Bendtsen *et al*., 2026), and the synthesis of transition state analogues, DUE-2,6-TDA and DUP-2,6-TDA, was described in detail in the supplementary information. These fragments were solubilised in 100% DMSO by vortexing several times to a final concentration of 1 M. For soaking, 1.0-1.5 µL of stock fragment solution was directly added onto a 6 µL crystal drop with a final concentration of compounds around 100 mM. Fragment soaking was performed from 10 min to 24 h. Crystals were then briefly soaked in 20-25 % glycerol mixed in mother liquor, fished out by nylon (Hampton Research) or LithoLoops (Molecular Dimensions) and flash frozen in liquid nitrogen. Compounds from the FragMAX fragment library (Lima *et al*., 2020) were solubilised in mother liquor to a final DMSO concentration of 10 % and briefly spun to remove air bubbles, after which 1-1.2 µL fragment buffer was directly added to crystal drops by multichannel pipette. For fishing and mounting fragments, the Easy Access Frame (Barthel *et al*., 2021) was used to prevent evaporation.

### 2.3. X-ray Diffraction Data collection, processing and structure solution

Initial crystal data collections were performed at the P13 and P14 beamlines at DESY, Hamburg and the I24 beamline at Diamond Light Source. Final X-ray diffraction data were collected on the BioMAX beamline at MAXIV (Lund, Sweden). All data sets were integrated and scaled by XDS (Kabsch, 2010). POINTLESS (Evans, 2011), was used to assign correct space groups. If auto-processed data from the autoPROC pipeline (Vonrhein et al., 2011) had reasonable statistics for high-resolution shell, it was used in the subsequent stages of structure determination, typically with a CC1/2 score above 0.3 was used for resolution cutoff. Initial templates for molecular replacement are generated by AlphaFold 3 (Abramson et al., 2024). Before molecular replacement, data sets were analysed by phenix.xtriage (Adams et al., 2010) and CTRUNCATE (Evans, 2011), to check for potential twinning or pseudo-translational issues. Molecular replacement was performed by Phaser (McCoy *et al*., 2007). Refinement and manual model building were performed primarily using phenix.refine (Afonine *et al*., 2012), REFMAC5 (Murshudov *et al*., 2011), and coot (Emsley *et al*., 2010), respectively. Ligand restraints were obtained by AceDRG (Long *et al*., 2017) and Elbow (Moriarty *et al*., 2009). Structure visualisation and figure preparation were performed in PyMOL (Schrödinger).

### 2.4. HPLC analysis of PU fragments hydrolysis

Endpoint samples were collected after incubation of 500 nM enzyme with 0.2 mg/mL DUE-TDA or DUE-MDA for 3 h at 40℃ in 100 mM bis-tris propane (pH 9.0) and 100 mM NaCl. The reaction was stopped by adding 100% acetonitrile at a 1:1 ratio, after which the aliquots were filtered through a 0.22 µm PTFE filter. All reactions were performed in duplicate. HPLC was used to quantify production of mono-substituted monourethane ethylene (MUE)-MDA and MUE-TDA, and their breakdown products 4,4-MDA and 2,4-TDA, respectively, using a Shimadzu system and UV-absorbance at 240 nm. 10 µL of sample was injected onto a Zorbax Eclipse Plus C18 column 5 µm particle size at 40 C and a flow rate of 1 mL/min. The mobile phase consisted of (A) milliQ water and (B) 100% acetonitrile. For DUE-TDA, a gradient separation was used from 30-80% B for 6 min, while DUE-MDA separation involved 5 min Isocratic elution (50% B). Standard curves of DUE-MDA, MUE-MDA, 4,4-MDA, DUE-TDA, MUE-TDA, and 2,4-TDA were used to assign peaks and convert areas to concentrations. All standard curves could be described by linear regression with R2 ≥ 0.95.

## 3. Results

### 3.1. Structural Characterisation of Amidase and Esterase

The determination of key positions and cavities for the catalytic properties of the enzymes requires comparison of the apo state with the complexed state. Therefore, we first determined the structure of the two enzymes u11 and AX^PUR5^in the ligand free state and then in the presence of a general suicide inhibitor, the latter helping to define the catalytic scaffold and the active site.

#### 3.1.1. Structural characterisation of amidase with and without a small suicide inhibitor reveals overall fold and catalytic pocket

The asymmetric unit in space group P 61 2 2 consists of a dimer of u11. A two-fold non-crystallographic symmetry axis relates the two monomers to the b-axis. In addition, we performed size-exclusion Chromatography to confirm a dimeric assembly in solution (**Fig. S1b**). The overall structure of the monomer consists of twelve α-helices (αA–L) and ten β-strands (β1–10), forming a twisted mixed β-sheet surrounded by alpha-helices on both sides, similar to an alpha–beta sandwich topology (**Fig. 2a**). The active site cavity is formed by four long loops (L1: 73–100, L2: 118–150, L3: 188–211, and L4: 348–375) and two α-helices (αI and αK) on one edge of the cavity. A non-crystallographic twofold symmetry axis is established by direct contact between αI and αJ helices, and the same two helices also form a channel connected to the catalytic pocket. The catalytic triad, composed of the nucleophilic Ser173, cisSer149, and Lys74, is located deep in the pocket and closer to αI and αK. The correct alignment of *ζNH*^+^_3_ of Lys74 in the catalytic triad is maintained by hydrogen bonding from Ser168, Ser150, and cisSer149 (**Fig. 2c**). Moreover, *cis*Ser also establishes two direct hydrogen bonds with the nucleophilic serine via the backbone amino group and Oγ due to the unusual *cis* configuration (**Fig. 2c**). A DALI search (Holm *et al*., 2023) found UMG-SP1-3, one of the first reported PU-degrading AS amidases (Branson *et al*., 2023), to be the closest structural homologue for u11 as it was expected since u11 was found through homology search to UMG-SP1(Rotilio *et al*., 2026). UMG-SPs share a very similar overall fold (i.e., Cα rmsd < 1.0 Å) and active site with u11, which indicates a highly conserved evolutionary catalytic scaffold. In summary, u11 represents a typical AS amidase catalytic scaffold that is highly conserved across bacteria, archaea, and eukaryotes (Chebrou *et al*., 1996).

**Figure 2.**
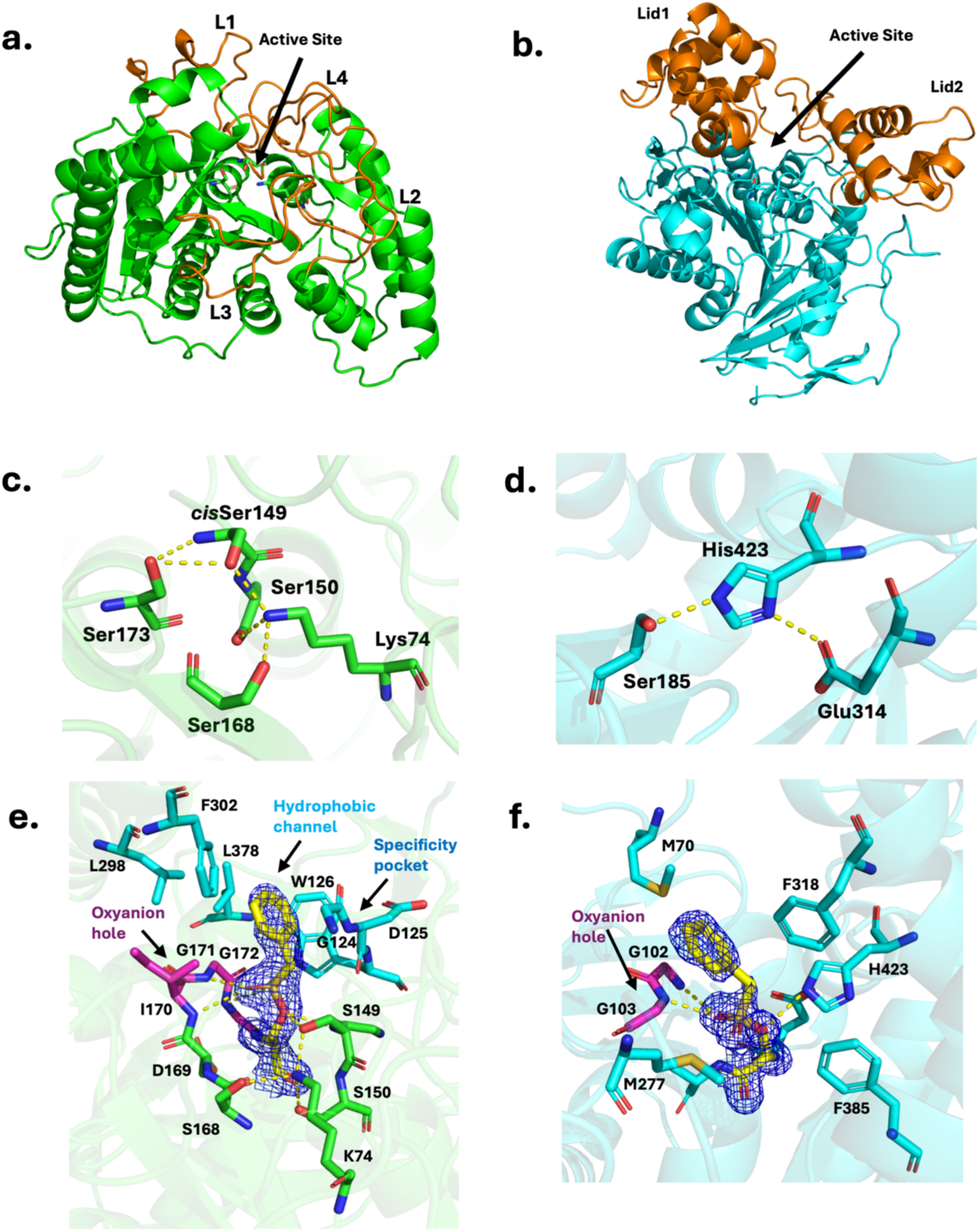
Structural overview of two Different Catalytic Scaffolds for Polyurethane Degradation. **a**) Overall structure of the amidase u11 with catalytic triad depicted as sticks and L1-4 shown in orange. b) Overall structure of the esterase AX^PUR5^shown with catalytic triad residues as sticks and Lid1 and Lid2 Domains shown in orange. Insets of the catalytic pentad of u11 c) and the catalytic triad of AX^PUR5^d). Structures of the PMS-bound region in u11. The 2mFo-DFc standard maps at 1.5 *σ* contour level for PMS is depicted in blue mesh e) and AX^PUR5^f) with surrounding residues shown as sticks and the 2mFo-DFc standard maps at 1.0 *σ* contour level for PMS is depicted in blue mesh.

To understand transition-state stabilisation and possible substrate recognition mechanisms, u11 was purified and crystallised in the presence of phenylmethylsulfonyl fluoride (PMSF), a catalytic mechanism-based suicide inhibitor commonly used against serine proteases. In the crystal structure, phenyl methyl sulfoxide (PMS) forms a covalent bond with the nucleophilic Ser173 (**Fig. 2e**), and a complete electron density map for PMS is available at 1.80 Å resolution. The sulfonyl group of PMS mimics a sp3-hybridised tetrahedral transition state. It is stabilised by the oxyanion hole formed by Gly171 and Gly172, whose backbone amine groups contact the oxygen atoms of the sulfonyl group. The benzyl group of PMS fits into the pocket formed by Trp126, Asp125, and Gly124, in which Trp126 stabilises a T-shaped π–π stacking interaction with the benzylic ring of PMS. Overall, this pocket resembles the specificity pockets also observed in several serine proteases that drive substrate selectivity (Hedstrom, 2002) and earlier described for signature amidases (Rotilio *et al*., 2025). Moreover, a hydrophobic channel formed by Leu298, Phe302, and Leu378 directly connects the bulk solvent to the catalytic pocket, constituting a possible route for substrate diffusion into the active site.

#### 3.1.2. Structural characterisation of esterase identified a TEV peptide ligand bound to the active site

The second enzyme, AX^PUR5^, is structurally and evolutionarily different from u11. AX^PUR5^ Crystallisation in the P 21 21 21 space group, with diffraction to 1.0 Å, revealing an asymmetric unit comprising a single polypeptide chain. Yet, gel filtration analysis suggested a dimeric state in solution (**Fig. S1a**). The monomeric structure of AX^PUR5^ is a centrally located alpha–beta hydrolase fold, with two alpha-helical lid domains atop the catalytic core (**Fig. 2b**). The active site is a deep cavity located between the two lid domains. It exhibits a highly hydrophobic character, surrounded by several aromatic residues, with the catalytic triad located at the centre (**Fig. 2d**). Nucleophilic Ser185, His423, and Glu314 comprise the catalytic triad, which is very similar to that of serine proteases (Di Cera, 2009). The catalytic serine is located on the nucleophilic elbow, as seen in numerous other carboxylic esterases and α/β hydrolases in general (Heikinheimo *et al*., 1999). In the ligand-free structure, the catalytic triad residues are connected by a hydrogen-bonding network that stabilises the catalytically active conformation. Strikingly, the ligand-free structure of AX^PUR5^ contains an unexpected peptide ligand captured between the two lid domains (**Fig. S2a**), whose sequence we can assign thanks to the near-atomic-resolution structure of the esterase, revealing leucine, tyrosine, and phenylalanine residues, as well as backbone density for other side chains. The LYF sequence is found in the TEV protease recognition sequence (-ENLYFQ-), used for His-tag cleavage in this construct, while residual density consistent with N– and Q-side chains further supports our hypothesis. Further refinement of the NLYFQ ligand yielded full density that clearly followed the peptide at a low contour level(**Fig S2a**), with a very strong omit map (**Table S2**). The Tyr and Phe side chain residues are inserted into a hydrophobic pocket lined by several aromatic residues. At the same time, the remaining part of the peptide is more solvent-exposed due to polar side chains. To our knowledge, this is the only structure in the Protein Data Bank (PDB) in which the TEV recognition peptide is bound to an enzyme other than TEV protease. Even though the peptide does not directly mimic plastic fragments, strong similarity to PU fragments in terms of size, bulkiness, and hydrophobicity implies probable binding of bulky plastic fragments at the same site.

To identify catalytically active residues, we purified in the presence of PMSF, a broad-spectrum serine hydrolase inhibitor. Indeed, in the resulting complex, PMS is covalently linked to the catalytic Ser185. At the same time, the sulfonyl oxygen interacts with backbone amine groups of the oxyanion hole (Gly102 and Gly103) via hydrogen bonding (**Fig. 2f**). Upon PMSF binding, the TEV peptide that we observed in the ligand-free structure cannot bind to a similar site because of the steric clash with benzylic rings of PMS and phenylalanine from the TEV peptide. There is no specific recognition pocket near the benzylic ring in AX^PUR5^, in contrast to the u11. Instead, the benzylic ring of PMS is fitted into the large hydrophobic pocket surrounded by aliphatic and aromatic side chain residues. The active site of the esterase is quite large and hydrophobic, allowing considerable substrate promiscuity.

### 3.2. Fragment-based active site probing maps potential subpockets in both enzymes

PMS-bound structures of the amidase and esterase enabled the characterisation of the catalytic pocket, highlighting the oxyanion hole and suggesting potential mechanisms for substrate recognition. Nevertheless, the existence of additional hydrophobic pockets distant from the catalytic pocket and potentially related to the binding of bulkier plastic fragments remains unclear. Therefore, we implemented a crystallographic fragment-screening campaign to probe the active-site cavity and map additional binding sites beyond the catalytic pocket. We used the FragMAX library (Lima *et al*., 2020), which consists of 160 high-concentration, high-solubility, non-redundant small molecules, on amidase and esterase crystals. As a result, we identified 9 hits clustering in two pockets for the amidase (e.g., aP1 and aP2) and 12 hits clustering in three pockets for the esterase (e.g., eP1, eP2 and eP3), with complete electron density for fragments in standard maps (**Fig. 3a** and **Fig. 4a**) and very strong omit maps for most of the fragments (**Fig. S3** and **Fig. S4**). Unfortunately, there is very little overlap in fragment binding between the two enzymes. Only one fragment (e.g., VT00167) is bound to both esterase and amidase (**Fig. S5**). An issue is that during soaking and data collection, many fragments induce crystal damage which results in low or no diffraction. This prevents systematic comparison of binding preferences for individual fragments to esterase and amidase.

**Figure 3.**
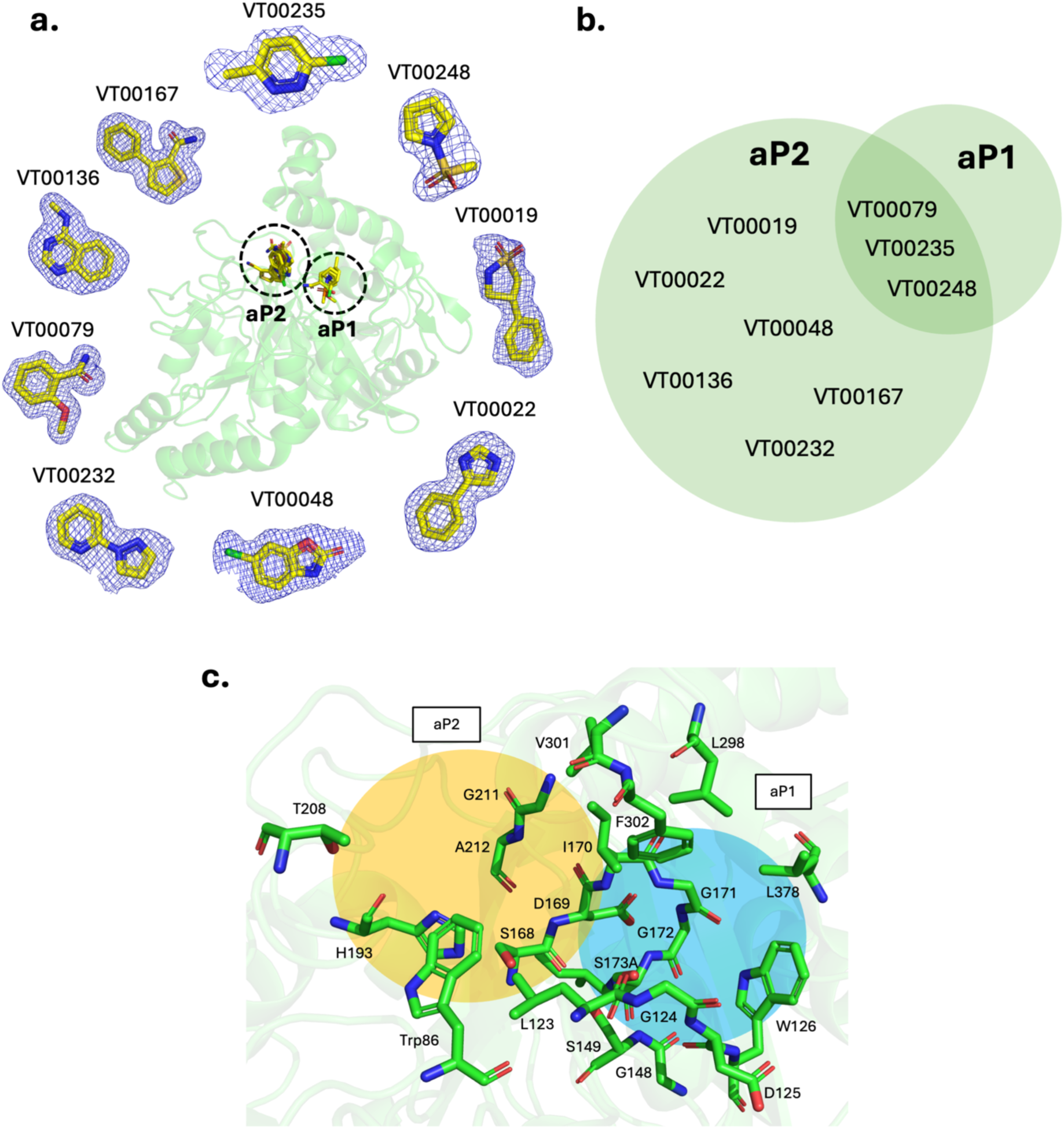
Fragment Based Active Site Mapping/Probing identifies potential pockets for plastic binding to esterase and amidase. **a**) Superimposed structures of fragments for amidase with each ligand demonstrated separately. The 2mFo-DFc standard maps at 1.5 *σ* contour level for Amidase fragments are shown in blue mesh. **b)** overlap of fragments that bound to aP1(3 hits) and aP2(9 hits) shown in circles with relative sizes corresponding to the number of hits. **c)** Close view of fragment binding pockets with surrounding residues highlighted, and the pockets are shown in shaded colour.

**Figure 4.**
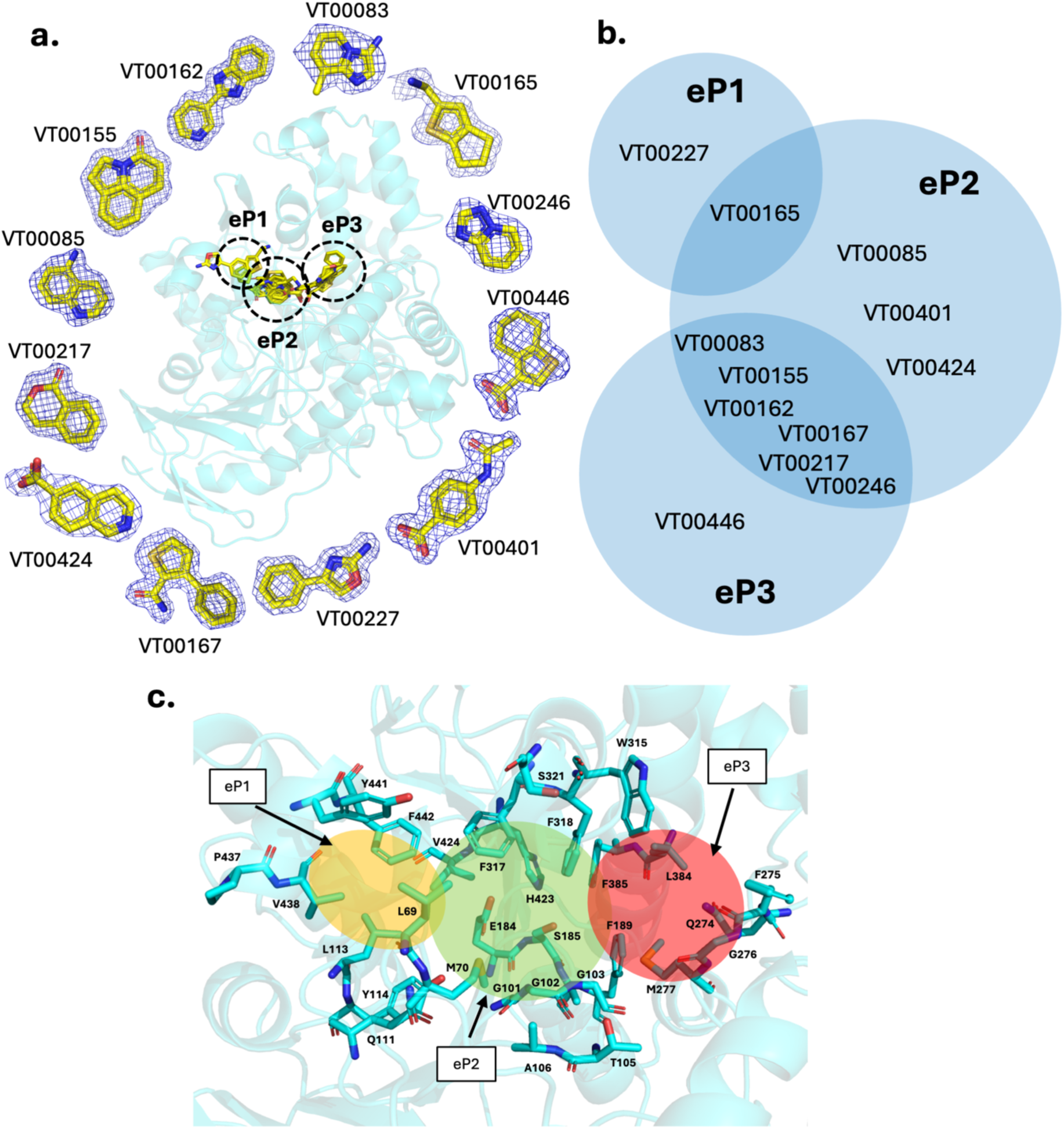
Fragment Based Active Site Mapping/Probing identifies potential pockets for plastic binding to esterase. **a**) Superimposed structures of fragments for esterase with each ligand demonstrated separately. The 2mFo-DFc standard maps at 1.0 *σ* contour level for esterase fragments is shown in blue mesh. **b)** The overlap of fragments that bound to eP1(2 hits), eP2(10 hits) and eP3(7 hits) is shown in circles with relative sizes corresponding to the number of hits. **c)** A close view of fragment binding pockets with surrounding residues highlighted and the pockets are shown in shaded colour.

#### 3.2.1. Amidase pocket mapping identifies two subpockets for potential PU binding

We identify two pockets in the amidase active site where aromatic fragment groups are clustered (**Fig. 3c**). A summary of the interactions of individual fragments with aP1 and aP2 is shown in **Table 1**. The first pocket (aP1) correlates with the catalytic pocket since it is the same place where PMS is covalently bound to the catalytic Ser173 and establishes interactions with the oxyanion hole (Ile170, Gly171, Gly172), catalytic pentad (Lys74, cisSer149, Ser150, Ser168, and Ser173), and substrate recognition pocket (Gly124, Asp125, Trp126) (**Fig. 3c**). Compounds VT00079, VT00235, and VT00248 bind to this site with a similar orientation of aromatic groups towards the specificity pocket (**Fig. 2e**). Fragment VT00079 bound to this pocket makes aromatic π–π interactions with Trp126 via its aromatic group, potentially constituting the molecular basis for substrate selectivity and recognition. Additionally, oxyanion hole backbone groups make hydrogen bonds with fragments if there is any donor/acceptor group available (**Table 1**), similar to the sulfonyl group interaction of PMS, which encapsulates TS stabilization (**Fig. 2e**). The aP1 is connected to the bulk solvent via a large hydrophobic channel at the top of the pocket that is surrounded by Leu298, Phe302, and Leu378 (**Fig. 3c**). Thus, it is highly probable that there are weak interactions of fragments in aP1 due to the solvent-exposed channel and weak hydrophobic interactions. Indeed, the hydrophobic channel that connects aP1 to the bulk solvent (**Fig. 2e**) might serve as a substrate/product binding/release route.

**Table 1.** List of fragments that map out binding pockets in the amidase.^1^.

| Fragment name | aP1 | Interacting residues at aP1 | Interaction type at aP1 | aP2 | Interacting residues at aP2 | Interaction type at aP2 |
| --- | --- | --- | --- | --- | --- | --- |
| VT00019 | - | - | - | + | Trp86, Ala122, Leu123, Ser149, Ser168, Asp169, Ile170, His198, Tyr200, Thr208, Val301, Phe302 | Hydrophobic interaction |
| VT00022 | - | - | - | + | Trp86, Leu123, Ser168, Asp169, Ile170, His198, Gly211, Ala212, Val301, Phe302 | Hydrophobic interaction |
| VT00048 | - | - | - | + | Trp86, Leu123, Ser168, Asp169, Ile170, His198, Thr208, Gly211, Val301, Phe302 | Hydrophobic interaction |
| VT00079 | + | Leu123, Gly124, Asp125, Trp126, Gly148, Ser149, Asp169, Ile170, Gly171, Gly172, Ala173, Phe302 | H-bonding, π-π interaction | + | Trp86, Ala122, Leu123, Ser168, Ile170, His198, Thr208, Val301, Phe302 | H-bonding, Hydrophobic interaction |
| VT00136 | - | - | - | + | Trp86, Ala122, Leu123, Ser168, Asp169, Ile170, His198, Thr208, Gly211, Ala212, Val301, Phe302 | Hydrophobic interaction |
| VT00167 | - | - | - | + | Trp86, Ala122, Leu123, Ser168, Asp169, Ile170, Ser195, His198, Thr208, Gly211, Ala212, Val213, Val301 | Hydrophobic interaction, π-π interaction |
| VT00232 | - | - | - | + | Trp86, Ala122, Leu123, Ser168, Asp169, Ile170, His198, Thr208, Gly211, Ala212, Val301, Phe302 | Hydrophobic interaction, π-π interaction |
| VT00235 | + | Gly124, Asp125, Trp126, Gly148, Ser149, Ile170, Gly171, Leu298, Phe302, Leu378 | Hydrophobic interaction | + | Trp86, Ala122, Leu123, Ser168, Asp169, Ile170, His198, Thr208, Gly211, Ala212, Val301, Phe302 | Hydrophobic interaction |
| VT00248 | + | Leu123, Gly124, Asp125, Trp126, Gly148, Ser149, Ile170, Gly171, Gly172, Ala173, Phe302 | Hydrophobic interaction, H-bonding | + | Trp86, Leu123, Ser168, Asp169, Ile170, His198, Gly211, Ala212, Phe302 | Hydrophobic interaction |

<sup>1</sup> Fragments that bind/do not bind to the pocket are indicated as + and -, respectively. Interacting residues within 4 Å and interaction types are written for corresponding pocket. Residues that are involved in H-bonding are highlighted in red, and residues that are involved in $\pi$ - $\pi$ interactions are highlighted in blue.

The second pocket (aP2) is located close to aP1, and is buried in the active site by Loop2 and Loop3. Notably, this pocket is deep, large, highly hydrophobic, and partially protected from the bulk solvent, in clear contrast to aP1 (**Fig. 3c**). All fragments bind to aP2 (**Fig. 3b**), indicating ligand promiscuity of this site compared to aP1. The aP2 is surrounded by Trp86, Leu123, Ser168, Ile170, His198, Gly211, Ala212, and Phe302. The major binding environment for ligands consists of hydrophobic interactions in addition to a few H-bonding and π–π interactions via Thr208 and Trp86, respectively. Aligned structures of ligand free structure and fragment-bound complexes indicate no significant conformational change in the surrounding residues, implying a lock–key-based enzyme–ligand interaction due to the large space available for bulky fragment binding(**Fig. S6**). Five– and six-membered aromatic ring groups of fragments occupy a similar position in the relatively large hydrophobic cavity of aP2 (**Fig. 3a**). In summary, aP2 generally binds more bulky and hydrophobic ligands as compared to the fragment found in aP1, which is more polar and exposed pocket with access to the solvent channel (Fig. 2e). Indeed, fragments with a single aromatic ring structure (i.e., VT00079, VT00235, and VT00248) can bind to aP1, whereas fragments with more than one ring structure cannot probe aP1 due to bulkiness and enhanced hydrophobicity (**Fig. 3a** and **Table 1**). This observation again supports the potentially important role of the aP2 site in binding more hydrophobic and bulky fragments that mimic PU aromatic groups.

#### 3.2.2. Esterase Pocket Mapping identifies three potential subpockets for PU binding

The fragment-based screening revealed more hits in AX^PUR5^ (12/160) than in u11, which we attribute to the larger and more solvent-accessible active site in the esterase. Three non-overlapping pockets (eP1, eP2, and eP3) are identified in the large active site cavity (**Fig. 4a**). **Table 2** summarises the fragments that bound to the corresponding pockets and the key interactions involved in those pockets. eP1 is located at the exit site of the large cavity, surrounded by Pro437, Val438, Tyr441, and Phe442, and binds two fragments (VT00165 and VT00227). Most of the fragments bind to eP2, which surrounds the nucleophilic Ser185. Specifically, fragments mostly interact with Ser185, the oxyanion hole (Gly102 and Gly103) and surrounding hydrophobic residues. Notably, fragments with H-bond donor/acceptor groups (i.e., VT00083, VT00085, VT00401, and VT00424) are oriented in a favourable conformation to establish hydrogen bonds with the side chains of Gln111, Tyr114, Glu184, Ser185, as well as backbone groups in the oxyanion hole (Gly102 and Gly103) (**Table 2**), thus mimicking possible interactions with the substrate and the leaving group (either amine or carboxyl) after urethane bond degradation.

**Table 2.**
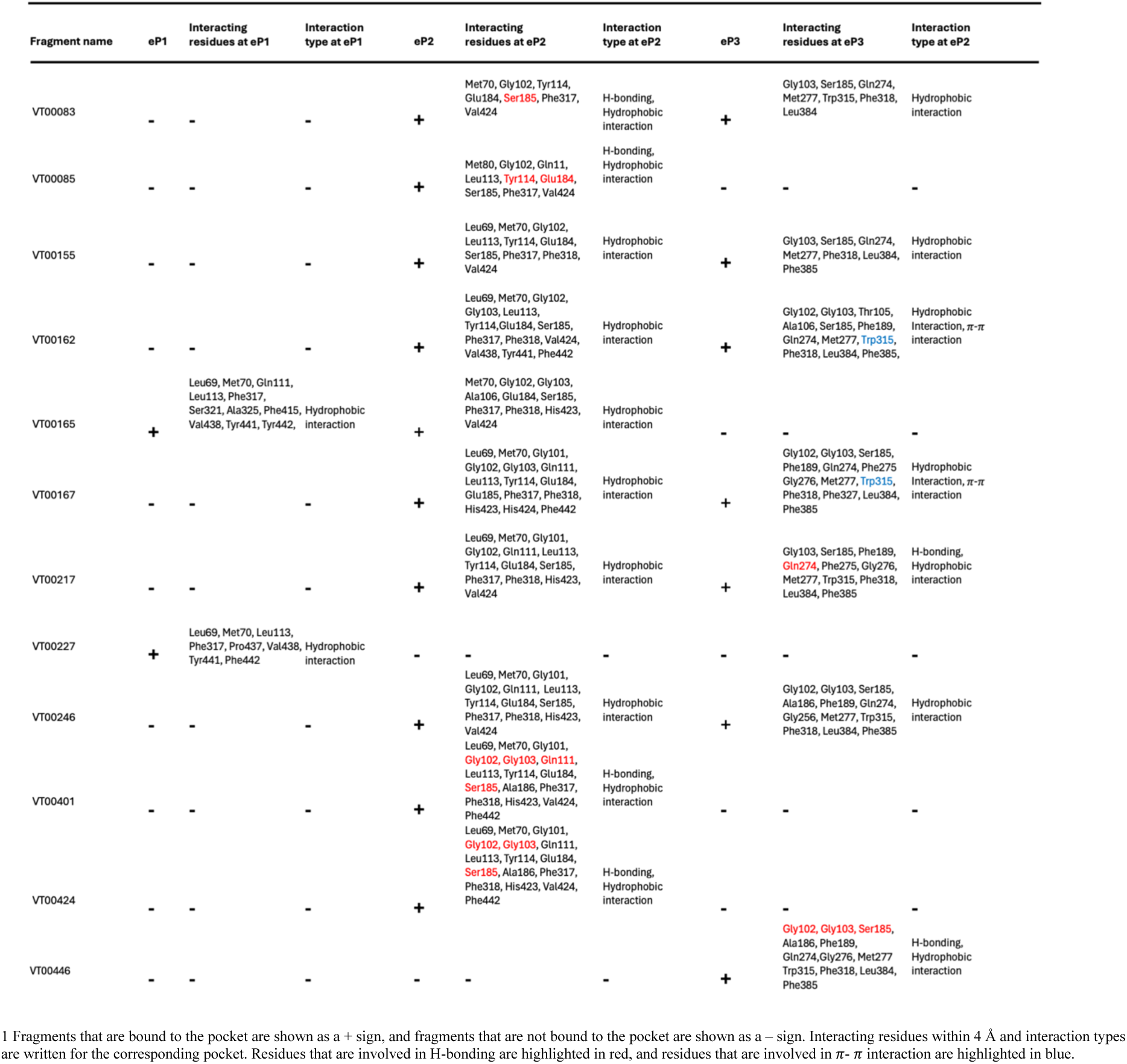
List of fragments that mapped the pockets in esterase.^1^.

eP3 is located slightly above eP2 and is buried deep within the hydrophobic cavity, partially overlapping the peptide-binding site (**Fig. 4b**). eP3 is surrounded by Phe189, Met277, Val384, and Phe385. Aromatic groups of the binding fragments (i.e., VT00162 and VT00167) partially overlap in this pocket, forming mostly hydrophobic interactions and a few favourable π–π stacking interactions with Trp315, as well as H-bonding interactions with Gly102, Gly103, Ser185, and Gln274 (**Table 2**). As observed in the ligand-free structure, the same site is protected from the solvent by a trapped TEV peptide fragment, in which Phe and Tyr rings of the TEV peptide extend towards this extremely hydrophobic pocket and might prevent the influx of an entropically unfavourable ordered solvent network.

All in all, the active site cavity of the esterase is highly hydrophobic and large, allowing binding of fragments that are bulkier and more hydrophobic compared to the amidase. Moreover, it is surrounded by several aromatic and aliphatic side chains that contribute to binding, as well as a few strategic polar side chains that can form hydrogen bonds and enhance specificity for the ligand. However, hydrophobic interactions provide the major force stabilising fragment binding (**Table 2**), again highlighting the hydrophobic nature of the active site of AX^PUR5^, appropriate for PU fragments. eP1 is targeted less by fragments and is more polar, whereas eP2 and eP3 are densely populated by hydrophobic side chains, which implies a potential role in binding aromatic groups of PU. Indeed, most fragments bind to eP2 and eP3 due to the larger, more hydrophobic space available to them, in contrast to the more solvent-exposed eP1. For example, bulky fragments like VT00155 and VT00162 bind to both eP2 and eP3 but not to eP1 (**Fig. 4a** and **Table 2**). Six fragments bind to both eP2 and eP3, whereas one fragment (VT00446) is unique to eP3 and three fragments (VT00085, VT00401, and VT00424) are unique to eP2 (**Fig. 4b**). This differential binding can be explained by two effects: i) preference for H-bonding with residues close to the given pocket and ii) differences in the local structure of fragments, especially branching of side chains from ring structures, which causes the fit of fragments in different pockets depending on the local stereochemistry (**Fig. 4c and Table 2**). In summary, the polar groups in eP1 and eP2 may help orient polar groups in PU and favour catalytically productive conformations, while eP2 and eP3 may bind bulky and hydrophobic ligands such as PU.

### 3.3. Validation of sub-pockets as PU fragment binding sites

The fragment-based pocket identification approach provides two and three pockets as potential binding sites for PU fragments in the amidase and esterase, respectively. To test whether any of these mapped pockets in AX^PUR5^ and u11 participate in the binding of bulky PU fragments, we used several PU analogues as TDA– and MDA-based substrates, intermediates, and product analogues. The main problem with hydrophobic plastic fragment soaking is the low solubility; many fragments precipitate when added to the crystal drop. We found it necessary to solubilise the fragments at high concentration (100 mM) in 100% DMSO stock to maximise solubility to saturate binding sites for low-affinity PU fragments for at least 2 h and up to > 20 h. This led to five different TDA-based polyurethane fragments bound to u11, in which the aromatic groups of all TDA fragments bind to aP2 (**Fig. 5**). For AX^PUR5^, we determined the structures of four plastic fragments, namely 4,4-MDA and DUE-TDA analogues, bound to eP1 and eP2 (**Fig. 6**).

**Figure 5.**
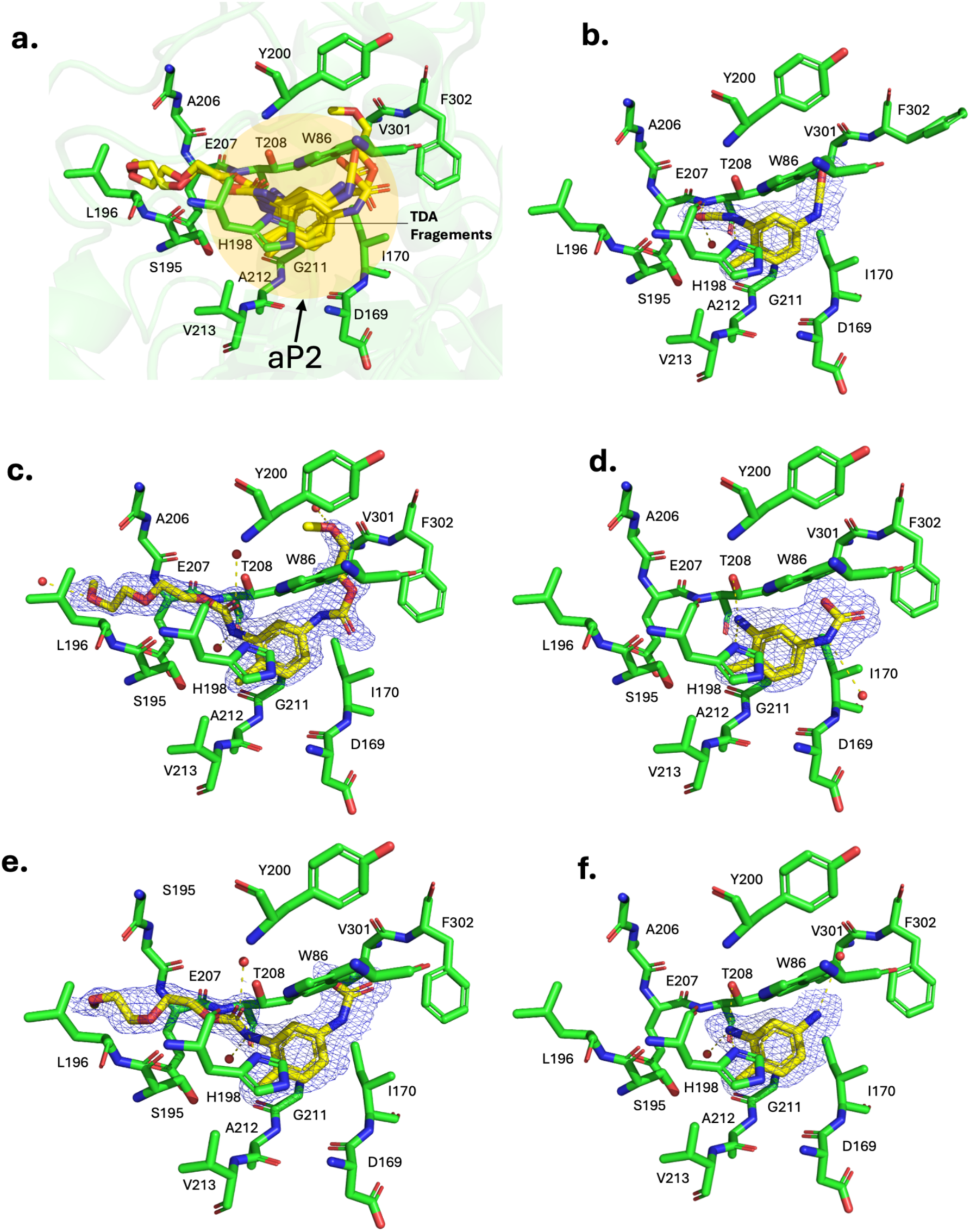
Validation of subpockets as plastic fragment binding sites in amidase. **a**) Superimposed structures of bound fragments in the aP2 and surrounding residue in the pocket are shown. b) 2,4-TDI c) DUE-2,4-TDA d) 2,4-TDA (carbamic acid) e) MUE-2,4-TDA (carbamic acid) and f) 2,4-TDA (amine) 2mFo-DFc map at 1.0 *σ* contour level in blue mesh for ligand along with surrounding residues. Hydrogen bonds are shown as dashed yellow lines and water molecules depicted as red spheres.

**Figure 6.**
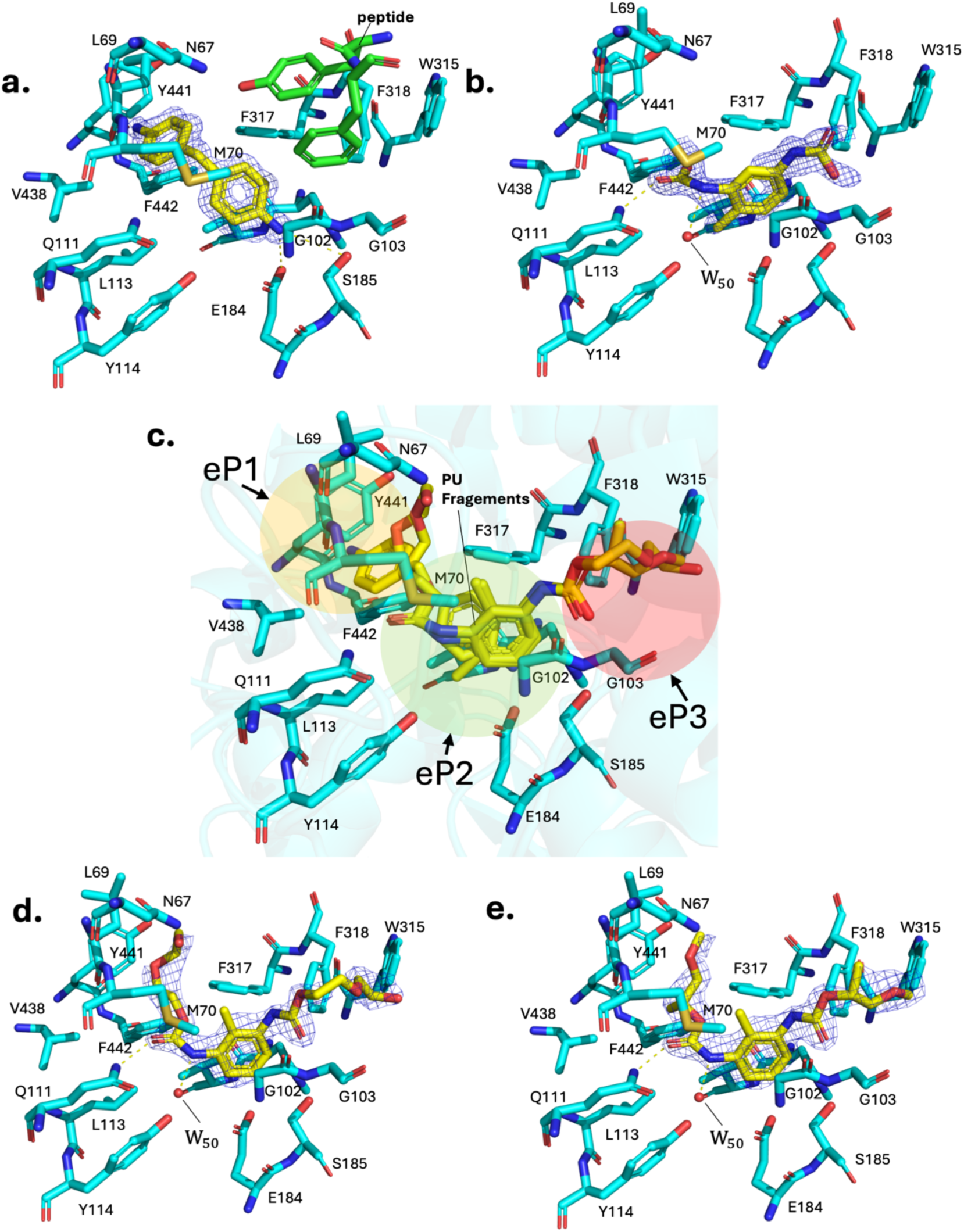
Validation of subpockets as plastic fragment binding sites in esterase. **a**) 4,4-MDA bound structure in the presence of trapped peptide (green) or b) 2,4-TDA (carbamate) degradation product, c) Superimposed structure of bound fragments in the eP1, eP2 and eP3 with surrounding residues are shown, d) 2,4-TDA (ethylene glycol) and e) 2,4-TDA (polypropylene). 2mFo-DFc map at 1.0 *σ* contour level in blue mesh is shown for fragments. Hydrogen bonds are shown as dashed yellow lines and water molecules depicted as red spheres.

#### 3.3.1. Amidase PU fragments bound structures show bulky TDA-based PU binding site at aP2

To obtain plastic fragments in the active site cavity, the inactive amidase mutant S173A was expressed, purified, and crystallised. For soaking experiments, various soluble plastic fragments, including DUE-MDA, DUE-2,4-TDA, MUE-MDA, MUE-2,4-TDA, 2,4-TDA, 2,4-TDI, and 4,4-MDA, were soaked in both u11-WT and u11-S173A. As a result, we captured both substrate and product analogues using the wild-type enzyme and the active-site mutant, which degraded substrates to different degrees (see below). Two TDA substrates, namely DUE-2,4-TDA (**Fig. 5c**) and MUE-2,4-TDA (**Fig. 5e**), and two degradation products, 2,4-TDA (amine) (**Fig. 5f**) and 2,4-TDA (carbamic acid) (**Fig. 5d**), and 2,4-TDI (substrate for TDA synthesis) (**Fig. 5b**), are obtained at high resolution, with visible density for all parts of the ligands in maximum-likelihood weighted 2mFo–DFc standard maps at 1.0σ contour level and no significant mFo–DFc difference map features at ±3σ contour level. Clear omit maps for ligands also confirm the binding (**Table S3**). Soaking of MDA-related compounds results in no clear density for ligands, even though partial/weak densities are observed at low contour levels in mFo–DFc, but no further modelling of ligands is performed.

All TDA fragments bind aP2, which was previously mapped by fragment screening as a potential binding site (**Fig. 3a**). Indeed, the binding of aromatic ring groups of fragments overlap with toluene diamine ring of TDA fragments (**Fig. S7f**.). Binding of long fragments displaces water molecules otherwise bound in the ligand-free structure (**Fig. S7b**). The toluene diamine ring is aligned almost perfectly in the same orientation in different TDA fragments in aP2 (**Fig. 5a**). Long ethylene glycol tails of DUE-2,4-TDA(**Fig. 5c**) and MUE-2,4-TDA(**Fig. 5e**) also bind and extend towards the same long tunnel in aP2, making several contacts with backbone groups of surrounding residues in the pocket. Geometrically, the tail segment fits very nicely into the pocket, indicating lock-and-key–based protein–ligand interaction, even though no significant hydrogen bonding contacts or salt bridges are observed. Soaking of DUE-2,4-TDA into wild type u11 crystals results in *in crystallo* degradation and capturing of 2,4-TDA carbamic acid. In the proposed reaction mechanism, urethane bond cleavage occurs via the release of an alcohol and a carbamic acid group, which subsequently hydrolyses spontaneously to an amine group (Paiva *et al*., 2025). However, the trapped TDA fragment is not directly exposed to bulk solvent and is therefore stabilised in a relatively hydrophobic cavity of aP2 that might prevent the spontaneous hydrolysis of the carbamic acid group of 2,4-TDA (**Fig.5d**). A similar carbamic acid moiety is also observed in the MUE-2,4-TDA–bound structure in the active site of mutant u11-S173A. This is most likely caused by residual activity remaining in the catalytic pocket by other serine residues in the hydrogen bonding network, as it has also been observed in serine proteases (Hedstrom, 2002). In summary, aP2, which is probed and mapped by fragment screening, indeed favours binding to TDA-based PU. Since both substrates and products bind to this pocket in a catalytically unfavourable orientation, aP2 is most likely related to binding and stabilising bulky TDA plastic to guide it towards the catalytic pocket. This observation confirms the potential role of aP2 in binding bulky, hydrophobic fragments, as shown by fragment-based pocket probing in the previous section. Moreover, reduced amidase activity towards TDA-based PU, compared with MDA-based PU (**Fig. S8b**), suggests potential product inhibition due to binding of TDA fragments, as the relative concentration of the product (e.g., MUE-TDA and 2,4-TDA) increases throughout catalysis and might remain bounded to aP2 which restrict the binding of the new substrate as we saw in the MUE-TDA (**Fig. 5e**), and 2,4-TDA(**Fig.5f**) bound structures.

#### 3.3.2. Esterase PU fragments bound structures show distinct modes of MDA and TDA based PU binding

Fragment soaking of PU fragments to AX^PUR5^ results in four high-resolution structures bound to 4,4-MDA, 2,4-TDA (carbamic acid), DUE-2,6-TDA, and DUP-2,6-TDA (**Fig. 6c**). All TDA-related fragments bind approximately at the same site, with slight differences in the flexible ethylene glycol arms. Toluene groups of fragments are located in the eP2, with arms extending towards eP3. The urethane bond in TDA fragments is near the nucleophilic serine, suggesting a plausible catalytic orientation (**Fig. 6bde**). On the other hand, 4,4-MDA, the fully degraded product for DUE-MDA, is slightly shifted from the TDA fragment location towards the eP1 location (**Fig. 6a**). The first benzylic ring of 4,4-MDA makes a direct hydrogen bond with the nucleophilic serine via its amine group whereas, the second benzylic ring of 4,4-MDA occupy the same location as fragment VT00227 (**Fig. S9d.**) which mapped the eP1. Considering the hydrogen-bonding distance, we captured the leaving group from the carbamate bond–degraded MDA-based polyurethane. Gln111 makes a hydrogen bond with the carbonyl oxygens of the urethane bond of all TDA-related fragments. A conserved water molecule (*i.e.*, water50) in the TDA binding pocket is found to interact with the amine group of the urethane bond of TDA fragments. This water molecule might be important for catalysis due to its proximity to the nucleophilic serine (**Fig. 6bde)**

Surprisingly, the PU fragments induce a conformational change in the peptide trapped between lid-1 and lid-2 (**Fig. S2**). Binding of 2,4-TDA releases the peptide fragment due to steric clash (**Fig. 5b**). In contrast, binding of DUE-2,6-TDA and DUP-2,6-TDA changes the peptide location and conformation completely, from the initial position (pose 1) to a more solvent-exposed position (pose 2) (**Fig. S10bd**). Both 2,6-TDA-related fragment structures are resolved in the presence of the peptide, forming what we term double-ligand structures, along with the 4,4-MDA-bound structure. The major difference between 2,4-TDA and 2,6-TDA-related fragments is the location of the methyl group (para in 2,4-TDA and ortho in 2,6-TDA) (**Fig. 1**). The exact reason for peptide release in 2,4-TDA and displacement in 2,6-TDA remains unclear. Still, it might relate to the initial binding mode of the fragments. On the other hand, 4,4-MDA binding does not alter the peptide’s spatial orientation or position, since the 4,4-MDA binding site is located away from the peptide binding site, implying no steric clash (**Fig. S10ac**). Moreover, when we soaked 4,4-MDA into PMSF-purified esterase crystals, we obtained a triple ligand complex in which 4,4-MDA, PMS, and peptide are bound to the esterase, although with reduced occupancy, as observed by a weaker but still positive unbiased omit map (mFo–DFc) at the 2.5σ contour level (**Fig. S10a**). We also noticed that the benzylic ring of PMS and the benzylic ring of phenylalanine from the peptide occupy the same location, consistent with partial occupancy of PMS and the peptide in different unit cells. There is no alternative conformation for phenylalanine from the peptide to occupy. Therefore, we modelled the peptide as NLYAQ by substituting phenylalanine with alanine. Even though the trapped peptide is not part of the structure, it acts as an extension of the loop in the Lid-1 domain, which contributes to the binding of plastic fragments by covering the active site and increasing overall hydrophobicity.

In summary, we obtained four high-resolution PU fragment–bound structures of AX^PUR5^: first with 2,4-TDA bound with displaced peptide; then three structures in the presence of 4,4-MDA and TDI fragments along with the peptide in different conformations (double ligand); and finally, in the presence of 4,4-MDA, PMS, and peptide (triple ligand). Comparison with fragments that are used to map binding sites reveals that aromatic moieties of fragments located at similar locations for eP1(i.e., VT00227) and eP2(i.e., VT00401) and makes similar type of interactions (**Fig. S9ef**). On the other hand, flexible tails of 2,6-TDA fragments partially overlap with aromatic groups of fragments (*i.e*., VT00446) at eP3 **(Fig. S9g).** All these structures imply extensive binding of PU fragments to the large active site of the esterase, in which fragments bind almost the entire available active site cavity. Since the active site of the esterase is larger and more solvent-exposed than that of amidase, the inherent flexibility of fragments is observed on urethane arms of 2,4-TDA and 2,6-TDA-related fragments, as evinced by electron density maps (**Fig. 6de**). We propose that the large and solvent-exposed active site cavity of esterase should favour the binding of bulky PU fragments as seen for double– and triple-ligand-bound structures. The 2,4-TDA-bound structure implies *in crystallo* degradation of DUE-TDA; however, this also leads to product inhibition, since 2,4-TDA should compete with the binding of substrate to the active site, as also observed in the lack of activity for DUE-TDA-based PU (Bendtsen *et al*., 2026). In addition, improper positioning of reactant groups of 2,6-TDA-based PU fragments due to the position of methyl group (**Fig. 1ef**) with respect to scissile urethane bonds indicates steric clash with active site residues and hence results in an unproductive and inhibitory binding pose as well. Finally, the orientation of 4,4-MDA suggests more favourable binding of MDA-based PU and catalysis by the esterase catalytic scaffold, as confirmed by the esterase’s demonstrated activity against MDA-based PU (Bendtsen *et al*., 2026).

### 3.4. Transition state analogues provide the catalytic snapshots of PU by extensively probing pockets

Finally, we turned to other types of inhibitors to obtain mechanistic details into the binding and stabilisation of catalytic intermediates and transition state analogues of PU. Inspired by PMSF, novel suicide inhibitors that mimic MUE-MDA and MUE-TDA were designed (Teixeira *et al*., 2026), which we synthesised and soaked into the two enzymes. Acylation of the substrate urethane bond and the subsequent tetrahedral intermediate are shown to be common steps in AS amidases (Cerqueira *et al*., 2017). Upon reaction with the nucleophilic serine, suicide inhibitors form covalent phosphoester bonds that mimic the hypothetical intermediate transition state during the first step of nucleophilic attack (acylation), in which serine hydrolyses the scissile carbonyl carbon (Teixeira *et al*., 2025; Paiva *et al*., 2025).

#### 3.4.1. MDA analogue inhibitor binds to two different conformations in the amidase active site

The MDA-TS inhibitor was soaked for different times, ranging from ten min to overnight. Analysis of the crystal structures clearly indicates that the formation of a covalent bond between the catalytic serine and the phosphate atom of the inhibitor takes place within 30 min. However, the appearance of clear density for the MDA ring structure and the flexible ethylene glycol tail requires a minimum of 1 h of soaking but complete density for ethylene glycol tails appeared after several hours (e.g., min 4 h)(**Fig. S11**).The large flexibility of the long ethylene glycol tail explains this behaviour, which is also reflected with clear residual densities in the difference map, even after several rounds of refinement without adding the ligand to the model. Strong density is observed near the catalytic serine at shorter times, which implies the presence of the phosphoryl group. This makes it evident that relaxation of the fragment, especially the ethylene glycol tail, and rotation of the methylene bridge in the MDA ring require more time to reach a stable conformation. Remarkably, our crystal structures also indicate that the inhibitor binds in two different conformations. In the first mode of binding (**Fig. 7a**), the ethylene glycol tail is oriented toward the aP1 and bent to fit into the space created by the flipping of Phe302. On the other side, the MDA ring fits nicely into the junction between the aP1 and the aP2, with a very good density in the 2mFo–DFc map, even at higher contour levels (1.5σ–1.7σ); however, the ethylene glycol tail has complete density at the 1σ contour level, as expected due to flexibility. In the second binding pose (**Fig. 7b**), the MDA ring is oriented into the aP1 and makes a T-shaped aromatic π–π stacking interaction with Trp126 through its first benzylic ring (**Fig. S7e**). The second benzylic ring of the inhibitor extends into the hydrophobic channel, as described in PMS (**Fig. 2e**), but shows weak/partial density, possibly because of flexibility or lack of clear interaction with nearby residues. The ethylene glycol tail in this second pose extends into the aP2 and has very good density for all parts, implying tighter interaction compared to the first binding mode. Altogether, the observation of two different binding modes for the MUE-MDA–based TS analogue indicates the versatility of the active site for binding to MDA-based PUR fragments in different orientations. Moreover, clear MDA-based PU degradation activity by the amidase (Rotilio *et al*., 2026) supports the mechanistic basis of promiscuous binding to MDA-based PU.

**Figure 7.**
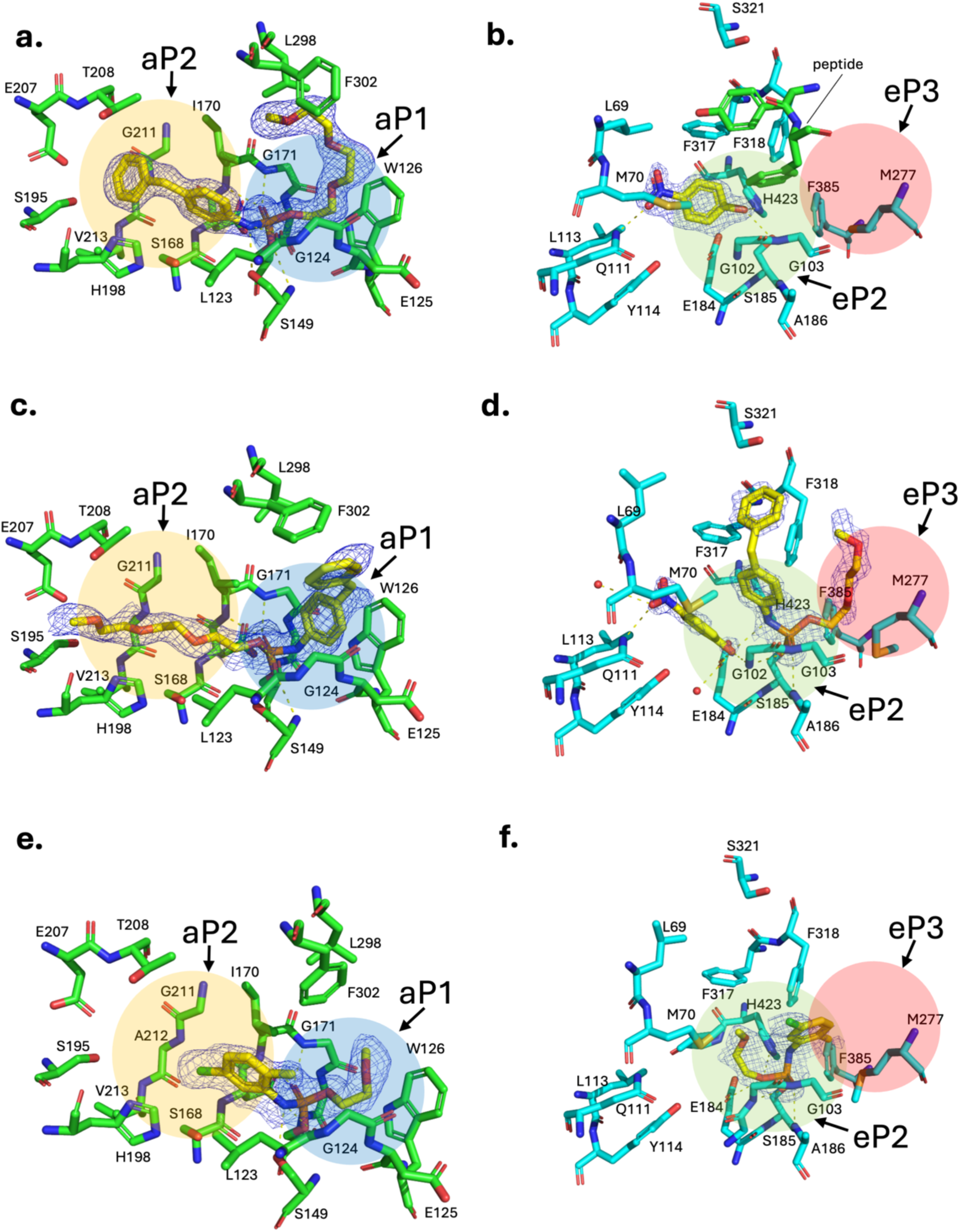
MUE-MDA and MUE-2,4-TDA Transition State analogues bound to amidase and esterase. **a**) u11 bound to MDA suicide inhibitor in pose-1. c) u11 bound to MDA suicide inhibitor in pose-2. e) u11 bound to TDA suicide inhibitor. b) AX^PUR5^ bound to leaving group p-nitrophenol d) AX^PUR5^bound to MDA suicide inhibitor. f) AX^PUR5^ bound to TDA suicide inhibitor. The 2mFo-DFc standard maps at 1.0 *σ* contour level are shown for fragments in blue mesh.

#### 3.4.2. TDA Analogue inhibitor binds to the same Pocket as TDA fragments in amidase

For the MUE-TDA TS analogue suicide inhibitor–bound crystal structure, we observed a similar tetrahedral covalent adduct in which a phosphoester bond is formed between O*_γ_* of Ser173 and the phosphate of the suicide inhibitor **(Fig. 7e).** Stabilisation of the double-bonded tetrahedral oxygen by the oxyanion hole is also observed, similarly to PMS and the MUE-MDA TS analogue. The aromatic group of the suicide inhibitor fits into the aP2, and the ethylene glycol tail extends toward the aP1. In comparison with the TDA fragment–bound structures, we see that the aromatic group of the suicide inhibitor is shifted toward the catalytic pocket. This state reproduces the catalytic snapshot during degradation, demonstrating that the TDA moiety would be trapped in the aP2 upon cleavage of the urethane bond, in clear agreement with our observations on other TDA fragment–bound structures. Further release of TDA from the aP2 and the later binding of a new TDA substrate would require loop motions. However, the additional ethylene glycol tail in the TDA group, which extends deep into the proximal pocket, hinders the release of product.

#### 3.4.3. MDA TS analogue and leaving group captured in esterase

In the proposed catalytic mechanism for carboxylesterases, a similar tetrahedral intermediate is formed upon nucleophilic attack of the catalytic serine on the carbonyl carbon of amide, ester, or carbamate bonds (Aranda *et al*., 2014). In this scenario, it could be expected that our suicide inhibitors would also bind to AX^PUR5^. To validate this theory, we first purified the esterase in the presence of the inhibitor, and furthermore, we co-crystallized and soaked for a long time in the first trial. Surprisingly, no structures contained a covalent bond between inhibitor and esterase; instead, we found the leaving group *para*-nitrophenol located close to the eP2 (**Fig. 7c**), establishing a hydrogen bond with the nucleophilic serine via its hydroxyl group, which resembles the fragments bound to eP2 (**Fig. 4a**). On the other hand, the nitro group formed another hydrogen bond with Asn111. Since *para*-nitrophenol bound tightly, we reasoned that an excess amount of *para*-nitrophenol remaining after synthesis and purification of the suicide inhibitor could remain bound to the active site. Thus, we synthesized a new batch of the inhibitor in which free *para*-nitrophenol is removed more thoroughly and tried soaking again. In this case, a covalent adduct between nucleophile Ser185 and the phosphor group formed. The ethylene glycol tail extends into the eP3 that is surrounded by several aromatic and aliphatic side chains, while the MDA core is on the other side of the pocket, where it is surrounded by several hydrophobic residues (**Fig. 7d**). Moreover, we also located the leaving group *para*-nitrophenol, which is released and captured just after the covalent bond is formed. Interestingly, one of the oxygens from the nitro group makes the same hydrogen-bonding interaction with Gln111 as we also observed in TDA fragment structures (**Fig. 6bde**). Moreover, the presence of the leaving group results in the upward conformation for the MDA core, which is almost orthogonal to the 4,4-MDA. The second benzylic ring of MDA faces towards Ser321, which might have made a potential hydrogen bond in the MUE-MDA bound structure (**Fig.7d).** It is highly probable that once the leaving group has been released, the MDA group fits into a similar site as the 4,4-MDA product. Additionally, a water molecule that is near to the hydroxyl group of para-nitrophenol (**Fig. 7d**) establishes a hydrogen bond similar to the conserved water molecule that we see in TDA fragments. Taken altogether, this water molecule is most likely the catalytic water molecule that attacks the urethane bond of TDA fragments and the tetrahedral TS.

#### 3.4.4. TDA TS analogue inhibitor binds in different pose than TDA fragments in esterase

The structure of AX^PUR5^ bound to a TDA analogue transition state inhibitor was solved at 1.4 Å (**Fig. 7f**). We expected to see a TDA group that fits into the same site (i.e., eP2 and eP3) as other TDA fragments (**Fig. 6c**). However, refinement of the TDA-bound suicide inhibitor was not straightforward, as we still observed positive difference peaks and a vague standard map associated with the fragments (**Fig.7f**) after several rounds of occupancy refinement. Nevertheless, the clear omit map for the fragment confirm the presence of TDA analogue transition state inhibitor(**Table S2**). This observation most likely reflects multiple conformations of the fragment. The lack of a single-state conformation for the ligand indicates weak stabilisation of the transition-state intermediate, which is essential for lowering the energy barrier for catalysis. We can see that the TDA group is located close to Met277 and Phe318, where the ethylene glycol tails of TDA bind. This binding pose for the toluene group differs from the 2,4-TDA fragment binding pose (**Fig. 6b**), which might also explain why the latter binding pose for 2,4-TDA-based PU is not suitable for catalysis. However, the lack of TDA degradation activity by the esterase (Bendtsen *et al*., 2026) suggests potential substrate inhibition, even though the first nucleophilic attack on the urethane bond yields a tetrahedral TS, as observed in this structure; subsequent hydrolysis of this intermediate would be unfavourable.

### 4.1. Discussion

In this study, we have demonstrated the efficiency of the Fragment-Based Active Site Exploration (FASE) approach to characterise interactions of different urethanases with various PU fragments. Our results highlighted several key features that could be essential to elucidate enzyme–plastic interaction at the molecular level. First, we mapped two binding pockets in the amidase (i.e., aP1 and aP2) and three pockets in the esterase (i.e., eP1, eP2 and eP3) in which our fragments are found clustered. Moreover, probing the active site of enzymes by fragments permitted the exploration of the binding landscape of novel enzymes for which real substrates are not known exactly. The simplicity of fragments that are used to map the active site effectively covers chemical space for potential sub pockets that can be important for substrate recognition. Additionally, we confirm the role of these probed sites for potential plastic fragment binding “hot spots” using a wide range of short, soluble plastic fragments that are analogues to the substrate, transition state, intermediate, and product. The mimicking of aromatic toluene and dianiline rings of PU by fragments clearly shows the applicability of the fragment-based approach as we probe similar types of interactions at mapped pockets (**Fig. S7 and Fig. S9**). By saturating binding pockets with fragment screening campaigns, we uncovered the important sites in active sites for PU binding and recognition. Additionally, by determining several structures of amidase and esterase in the presence of various PU fragments, we demonstrate the distinct binding modes of MDA– and TDA-based PU to completely different catalytic scaffolds and uncover the mechanistic level of differential activities of enzymes towards MDA– and TDA-based plastics. As a result, we demonstrated that by probing the active site with small-molecule compounds, we could map the actual plastic-binding sites that are in catalytically favourable orientations and positions within the large active-site cavity of our target enzymes.

### 4.2. Extensive PU binding by the entire active site of esterase

In the esterase, different binding modes of MDA– and TDA-based PU fragments were observed due to the large hydrophobic active-site cleft, which exhibits a high degree of substrate promiscuity, as shown in our fragment screening, plastic ligands, and trapped peptide fragment–bound structures. Depending on the active site shape, MDA and TDA cores are positioned differently, either in catalytically favourable or inhibitory poses. By targeting the surrounding residues around the TS analogues, it is possible to stabilize and hence reduce the activation energy that is required for catalysis or facilitate the release of the leaving group by weakening the interaction between the product and eP1 residues. Noteworthy, the capturing of *para*-nitrophenol and 4,4-MDA at the same binding site highlights the location for the product release route, which can be further engineered to reduce steric hindrance and weaken favourable interactions with the leaving group. On the other hand, all TDA fragments bind to the eP2 with almost the same orientation of the aromatic moiety and urethane bonds. In contrast, flexible ethylene glycol tails sample multiple conformations, as it evinced by weakened electron density maps towards eP3 (**Fig. 6bde**). Despite the favourable position of urethane bonds of TDA fragments for catalysis, the presence of a methyl group in the toluene ring prevents the proximity between the nucleophilic serine and the urethane bond for actual catalysis, thus correlating with the reported residual activity of this esterase for TDA-based plastic (Bendtsen *et al*., 2026). There are a few carboxylic esterases that are reported to degrade PU (Bendtsen *et al*., 2026). Interestingly, we only find one homologous esterase in which the structure is deposited in the PDB (8XRZ) along with an MDA degradation product–like MDA analogue suicide inhibitor that we used in this study. The aniline moiety of the MDA degradation product is located in a site analogous to the eP3 and the peptide binding site that we show in our structures. In contrast, MDA fragments that we observed in our structures (**Fig. 6a** and **Fig. 7d**) are located in a different site than in 8XRZ, which could be associated to local residue differences in eP3 and the peptide binding site. Nevertheless, this observation further enhances our conclusion, as we expected to see bulky PU fragments in the eP3 and the peptide binding site. Altogether, for a catalytically favourable conformation, the MDA group favours orientation towards the eP1, whereas TDA groups are buried toward the eP2 that is mapped by fragment screening. Moreover, the large hydrophobic active site of AX^PUR5^ offers the opportunity for substrate promiscuity through multiple binding conformations, although some binding poses may be unproductive or inhibitory.

### 4.3. Binding and guidance of bulky PU fragment by aP2 for catalysis in amidase

In the case of the amidase u11, aP2, which was mapped by our fragment screening, clearly stabilises the aromatic moiety of TS analogues and binds the substrate and product analogues of TDA. Moreover, in the homologous urethanase UMG-SP2, the same pocket binds to 4-hydroxybutyl(4-(4-aminobenzyl)phenyl) carbamate (BBC), which is a short-chain carbamate (Li *et al*., 2025). Even though BBC is not a true mimetic for either the toluene or aniline ring of PU, it further supports our observations that aP2 is important for binding bulky fragments. The relevance of the aP1 and the nearby specificity pocket, which bind to small aromatic carbamates in a favourable conformation for catalysis and product release, is highlighted by the MDA TS analogue–bound structure in this study (**Fig. 7c**), the degradation product 4,4-MDA–bound structure in a homologous urethanase (Rotilio *et al*., 2026), and QM/MM calculations on UMG-SP2 (Swiderek *et al*., 2025). In other AS amidases, such as FAAH (Min *et al*., 2019) and malonamidase (Shin *et al*., 2003), analogous specificity pockets and surrounding binding sites are observed, with differences in polarity and shape that enable binding to different types of substrates. This observation again highlights the specificity required for catalytically productive PU fragment binding. There are two distinct binding poses for the catalysis of the urethane bond in MDA-and TDA-based PUs. The orientation of the aromatic moiety of both TDA and MDA can either fall into aP2 (**Fig. 5a**) or extend towards the aP1 and substrate release channel (**Fig. 2e**), in the case of MDA. In the first scenario, loop dynamics are required to release the product, suggesting that further optimisation in loops could increase the catalytic turnover by enhancing flexibility. Indeed, engineering studies of homologous enzymes show that mutating residues on the same loop that covers aP2 significantly improves catalytic activity (Bayer *et al*., 2025; Li *et al*., 2025; Rotilio *et al*., 2025). In the second scenario, at aP1 the specificity pocket (**Fig. 2e)** interacts with and stabilises the first benzylic ring of the MDA moiety, as demonstrated in this study by PMS and described in Rotilio *et al*., (2025) **(Fig. 2e)** and the MDA-TS analogue inhibitor **(Fig. 7c)** and in previous docking and MD simulations (Paiva *et al*., 2025; Zheng *et al*., 2026). However, for TDA-based PU, the aP1 cannot bind to the toluene ring due to steric clashes with the amine and methyl groups. Furthermore, fragment screening also indicates that all fragments bind to aP2, whereas only three fragments with a single aromatic ring probe the aP1 (**Fig. 3a** and **Table 1**). However, aP2 needs to be carefully investigated for engineering since the same site also binds various TDA fragments, including substrate and product, in which extensive stabilisation of aromatic groups of PU fragments by engineering might lead to product inhibition and reduce catalytic turnover. Release of the product can be facilitated by engineering the loop that covers the aP2 by improving flexibility and reducing hydrophobic interactions, which again emphasises the important role of loops in ligand binding and turnover. In summary, aP2 is important for binding and stabilising long, bulky PU fragments for catalysis, whereas the aromatic groups of PU fragments need to bind to aP1 and extend into the hydrophobic channel to favour favourable product-release orientation and catalysis.

### 4.4. Structural features of different catalytic scaffolds for PU binding and degradation

The fragment-based approach for urethanases identifies several important features for binding and recognition of PU fragments **(Fig. 8).** For the esterase AX^PUR5^, we unravelled key structural features as: 1) a product release route in eP1; 2) promiscuous PU binding sites that spanning eP2 and eP3; 3) substrate access by large hydrophobic cleft; 4) Potential binding of bulky PU fragments at the peptide pocket and eP3 **(Fig. 8a).** For the amidase u11, the study revealed key structural elements, including: 1) a substrate and product diffusion channel; 2) binding of bulky PU fragments and their guidance towards catalytic site by aP2; 3) substrate capping by Phe302; 4) distinct orientations of TDA and MDA based PU for catalytically favourable conformation **(Fig. 8b).**

**Figure 8:**
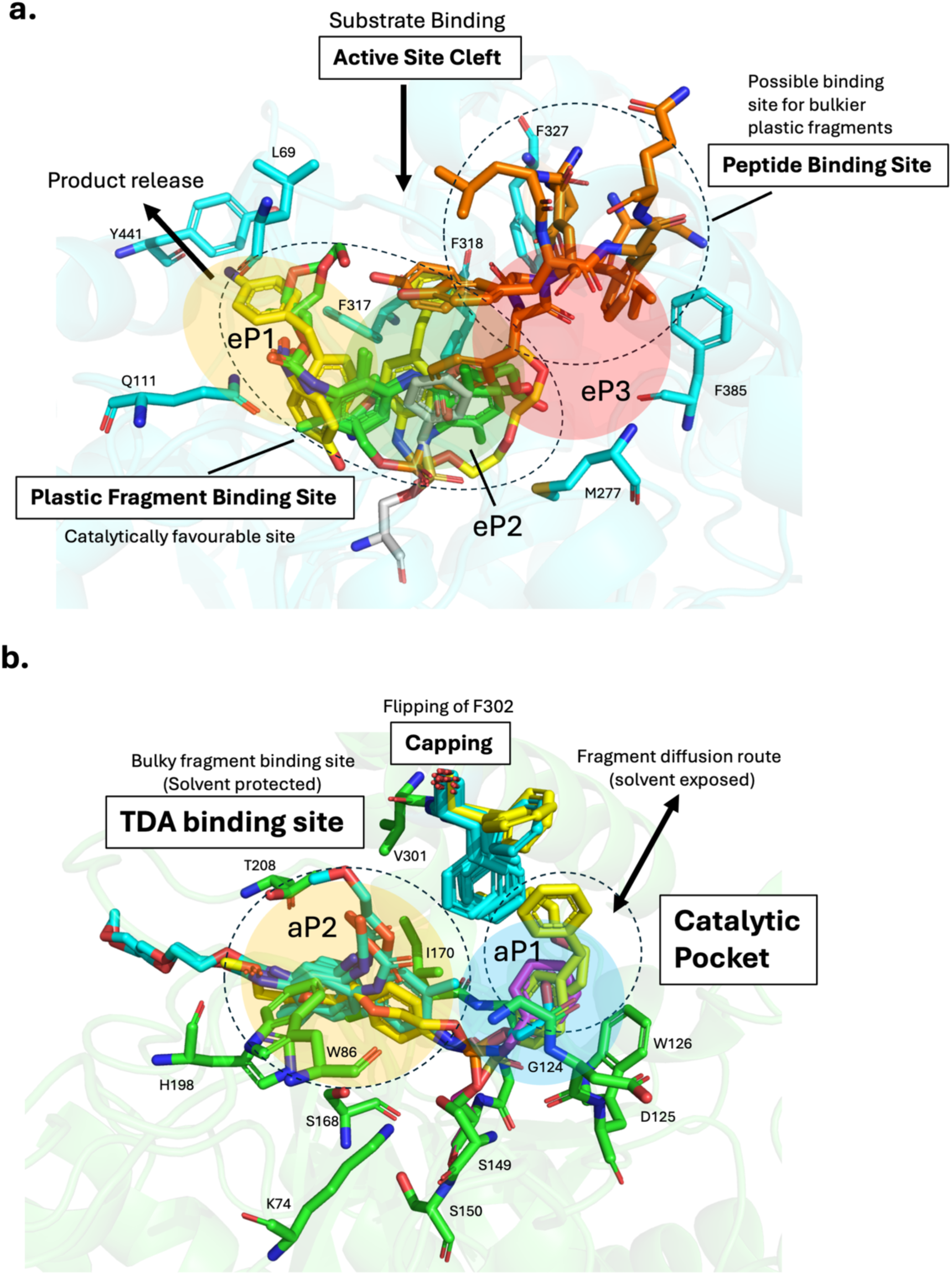
Summary of structural features for amidase and esterase identified by the FASE approach. **a**) Esterase structural features for plastic fragment binding. b) Amidase structural features. Mapped pockets are shown as shaded colour, and important binding sites are shown as dashed circles.

The fragment-based approach elucidates the molecular mechanism of interaction between plastic fragments and two different catalytic scaffolds that degrade polyurethane. This mechanistic understanding of the specificity for TDA and MDA based PU by different catalytic scaffolds and active sites holds the potential to guide engineering and discovery campaigns for homologous enzymes efficiently, since the overall structure of active sites and binding pockets are highly conserved in several reported enzymes (Chen *et al*., 2025; Bendtsen *et al*., 2026; Rotilio *et al*., 2026). Future studies can benefit from our fragment-based approach to identify “hot-spots” for engineering campaigns in homologous enzymes.

## 5. Acknowledgements

EnZync is funded by challenge grant NNF22OC0072891 from the Novo Nordisk Foundation. D.K. is financially supported by Erasmus+ program. We thank Tobias Krojer and Afshan Begum for assisting with FragMAX library soaking experiments. We thank beamline scientists Ana Gonzales, Isabel Bento, and Ezequiel Panepucci for assisting with data collection from the BioMAX beamline in MAXIV from proposals 20241360, 20250330, and 20251351. We thank Sofia Zhao for assisting data collection from I24 beamline from Diamond Light Source from proposal MX36082 for the AX^PUR5^ligand free and PMS bound state. We thank David von Stetten for assisting with data collection from the EMBL-P14 at PETRA-III in DESY for the initial characterisation of the u11 crystal from proposal MX992. Finally, we thank all member of EnZync.

## 6. Author Contribution

D.B. designed and conducted all experiments, analysed all data and prepared the manuscript. D.K. assisted with the expression and purification of proteins. L.R. provided the plasmids of u11 and u11-S173A. R.G. assisted with HPLC analysis. S.P.D expressed and purified the u11 for HPLC analysis. N.L.V. synthesised and purified suicide analogue inhibitors. A.S. and M.B.J. synthesised and purified PU fragments. S.S.T, J.P.M and D.E.O. supervised the project. D.E.O. and J.P.M. were responsible for funding acquisition and project management. All authors read and reviewed the final version of the manuscript.

## References

1. Abramson, J., Adler, J., Dunger, J., Evans, R., Green, T., Pritzel, A., … & Jumper, J. M. (2024). Accurate structure prediction of biomolecular interactions with AlphaFold 3. Nature, 630(8016), 493–500.

2. Adams, P. D., Afonine, P. V., Bunkóczi, G., Chen, V. B., Davis, I. W., Echols, N., … & Zwart, P. H. (2010). PHENIX: a comprehensive Python-based system for macromolecular structure solution. Biological crystallography, 66(2), 213–221.

3. Afonine, P. V., Grosse-Kunstleve, R. W., Echols, N., Headd, J. J., Moriarty, N. W., Mustyakimov, M., … & Adams, P. D. (2012). Towards automated crystallographic structure refinement with phenix. refine. Biological crystallography, 68(4), 352–367.

4. Akindoyo, J. O., Beg, M. H., Ghazali, S., Islam, M. R., Jeyaratnam, N., & Yuvaraj, A. R. (2016). Polyurethane types, synthesis and applications–a review. Rsc Advances, 6(115), 114453–114482.

5. Aranda, J., Cerqueira, N. M. F. S. A., Fernandes, P. A., Roca, M., Tuñón, I., & Ramos, M. J. (2014). The catalytic mechanism of carboxylesterases: a computational study. Biochemistry, 53(36), 5820–5829.

6. Banik, J., Chakraborty, D., Rizwan, M., Shaik, A. H., & Chandan, M. R. (2023). Review on disposal, recycling and management of waste polyurethane foams: a way ahead. Waste Management & Research, 41(6), 1063–1080.

7. Barthel, T., Huschmann, F. U., Wallacher, D., Feiler, C. G., Klebe, G., Weiss, M. S., & Wollenhaupt, J. (2021). Facilitated crystal handling using a simple device for evaporation reduction in microtiter plates. Applied Crystallography, 54(1), 376–382.

8. Bayer, T., Palm, G. J., Berndt, L., Meinert, H., Branson, Y., Schmidt, L., … & Bornscheuer, U. T. (2024). Structural elucidation of a metagenomic urethanase and its engineering towards enhanced hydrolysis profiles. Angewandte Chemie International Edition, 63(38), e202404492.

9. Bendtsen, M. K., Møllebjerg, A., Peña-Díaz, S., Graham, R., Petersen, N. C., Isaksen, B. N., … & Otzen, D. E. (2026). Environmental Identification of Novel Enzymes for Polyurethane and Polyamide Degradation. Angewandte Chemie International Edition, e6159643.

10. Branson, Y., Söltl, S., Buchmann, C., Wei, R., Schaffert, L., Badenhorst, C. P., … & Bornscheuer, U. T. (2023). Urethanasen für die enzymatische Hydrolyse niedermolekularer Carbamate und das Recycling von Polyurethanen. Angewandte Chemie, 135(9), e202216220.

11. Cerqueira, N. M. F. S. A., Moorthy, H., Fernandes, P. A., & Ramos, M. J. (2017). The mechanism of the Ser-(cis) Ser-Lys catalytic triad of peptide amidases. Physical Chemistry Chemical Physics, 19(19), 12343–12354.

12. Chebrou, H., Bigey, F., Arnaud, A., & Galzy, P. (1996). Study of the amidase signature group. Biochimica et Biophysica Acta (BBA)-Protein Structure and Molecular Enzymology, 1298(2), 285–293.

13. Chen, Y., Sun, J., Shi, K., Zhu, T., Li, R., Li, R., … & Wu, B. (2025). Glycolysis-compatible urethanases for polyurethane recycling. Science, 390(6772), 503–509.

14. Di Cera, E. (2009). Serine proteases. IUBMB life, 61(5), 510–515.

15. Emsley, P., Lohkamp, B., Scott, W. G., & Cowtan, K. (2010). Features and development of Coot. Biological crystallography, 66(4), 486–501.

16. Evans, P. R. (2011). An introduction to data reduction: space-group determination, scaling and intensity statistics. Biological crystallography, 67(4), 282–292.

17. Hedstrom, L. (2002). Serine protease mechanism and specificity. Chemical reviews, 102(12), 4501–4524.

18. Heikinheimo, P., Goldman, A., Jeffries, C., & Ollis, D. L. (1999). Of barn owls and bankers: a lush variety of α/β hydrolases. Structure, 7(6), R141–R146.

19. Holm, L., Laiho, A., Törönen, P., & Salgado, M. (2023). DALI shines a light on remote homologs: One hundred discoveries. Protein Science, 32(1), e4519.

20. Jhoti, H., Williams, G., Rees, D. C., & Murray, C. W. (2013). The’rule of three’for fragment-based drug discovery: where are we now?. Nature reviews Drug discovery, 12(8), 644–644.

21. Kabsch, W. (2010). xds. Biological crystallography, 66(2), 125–132.

22. Li, Z., Han, X., Cong, L., Singh, P., Paiva, P., Branson, Y., … & Liu, W. (2025). Structure-Guided Engineering of a Versatile Urethanase Improves Its Polyurethane Depolymerization Activity. Advanced Science, 12(13), 2416019.

23. Lima, G. M., Talibov, V. O., Jagudin, E., Sele, C., Nyblom, M., Knecht, W., … & Mueller, U. (2020). FragMAX: the fragment-screening platform at the MAX IV Laboratory. Biological Crystallography, 76(8), 771–777.

24. Long, F., Nicholls, R. A., Emsley, P., Gražulis, S., Merkys, A., Vaitkus, A., & Murshudov, G. N. (2017). AceDRG: a stereochemical description generator for ligands. Biological Crystallography, 73(2), 112–122.

25. McCoy, A. J., Grosse-Kunstleve, R. W., Adams, P. D., Winn, M. D., Storoni, L. C., & Read, R. J. (2007). Phaser crystallographic software. Applied Crystallography, 40(4), 658–674.

26. Min, C. A., Yun, J. S., Choi, E. H., Hwang, U. W., Cho, D. H., Yoon, J. H., & Chang, J. H. (2019). Comparison of Candida albicans fatty acid amide hydrolase structure with homologous amidase signature family enzymes. Crystals, 9(9), 472.

27. Moriarty, N. W., Grosse-Kunstleve, R. W., & Adams, P. D. (2009). electronic Ligand Builder and Optimization Workbench (eLBOW): a tool for ligand coordinate and restraint generation. Biological Crystallography, 65(10), 1074–1080.

28. Murshudov, G. N., Skubák, P., Lebedev, A. A., Pannu, N. S., Steiner, R. A., Nicholls, R. A., … & Vagin, A. A. (2011). REFMAC5 for the refinement of macromolecular crystal structures. Biological crystallography, 67(4), 355–367.

29. Teixeira, L. M., Paiva, P., Ferreira, P., Rotilio, L., Morth, J. P., Otzen, D. E., … & Ramos, M. J. (2025). Bridging experiment and theory: a computational exploration of UMG-SP3 dynamics. Pure and Applied Chemistry, 97(10), 1405–1418.

30. Teixeira, L. M., Paiva, P., Rotilio, L., Preben Morth, J., Fernandes, P. A., & Ramos, M. J.(2026). Leveraging Mechanism-Based Inhibition to Enable the Engineering of Enzymatic Activity towards Polyurethane. ChemRxiv. 2026–02.

31. Paiva, P., Teixeira, L. M. C., Wei, R., Liu, W., Weber, G., Morth, J. P., … & Ramos, M. J. (2025). Unveiling the enzymatic pathway of UMG-SP2 urethanase: insights into polyurethane degradation at the atomic level. Chemical Science, 16(5), 2437–2452.

32. Polecki, K., Paciorek-Sadowska, J., Borowicz, M., Isbrandt, M., & Zarzyka, I. (2026). Polyurethane Recycling: Sustainable Development Perspectives and Innovative Approaches. Materials, 19(4), 805.

33. Rossignolo, G., Malucelli, G., & Lorenzetti, A. (2024). Recycling of polyurethanes: where we are and where we are going. Green Chemistry, 26(3), 1132–1152.

34. Rotilio, L., Bayer, T., Meinert, H., Teixeira, L. M., Johansen, M. B., Sommerfeldt, A., … & Morth, J. P. (2025). Structural and functional characterization of an amidase targeting a polyurethane for sustainable recycling. Angewandte Chemie International Edition, 64(7), e202419535.

35. Rotilio, L., Østergaard, R. R., Thiesen, E. M., Paiva, P., Dwyer, K. M., Johansen, M. B., … & Morth, J. P. (2026). Expanding the Enzymatic Landscape for Polyurethane Degradation of Novel Bacterial Urethanases. bioRxiv, 2026-02.

36. Scott, D. E., Coyne, A. G., Hudson, S. A., & Abell, C. (2012). Fragment-based approaches in drug discovery and chemical biology. Biochemistry, 51(25), 4990–5003.

37. Shin, S., Yun, Y. S., Koo, H. M., Kim, Y. S., Choi, K. Y., & Oh, B. H. (2003). Characterization of a novel Ser-cisSer-Lys catalytic triad in comparison with the classical Ser-His-Asp triad. Journal of Biological Chemistry, 278(27), 24937–24943.

38. Swiderek, K., Arafet, K., de Sousa Batista, V., Grajales-Hernández, D., López-Gallego, F., & Moliner, V. (2025). Insights into the Catalytic Activity of a Metagenome-Derived Urethanase. Journal of the American Chemical Society, 147(46), 42511–42523.

39. Vonrhein, C., Flensburg, C., Keller, P., Sharff, A., Smart, O., Paciorek, W., … & Bricogne, G. (2011). Data processing and analysis with the autoPROC toolbox. Biological crystallography, 67(4), 293–302.

40. Zheng, M., Liu, J., Zhu, X., Chen, J., Zhang, Q., Wang, W., … & Li, Y. (2026). Computational Redesign of a Urethanase for Efficient Polyurethane Depolymerization. ACS Catalysis, 16(7), 6577–6588.

